# DISC1 regulates N-Methyl-D-Aspartate receptor dynamics: Abnormalities induced by a *Disc1* mutation modelling a translocation linked to major mental illness

**DOI:** 10.1101/349365

**Authors:** Elise L.V. Malavasi, Kyriakos D. Economides, Ellen Grünewald, Paraskevi Makedonopoulou, Philippe Gautier, Shaun Mackie, Laura C. Murphy, Hannah Murdoch, Darragh Crummie, Fumiaki Ogawa, Daniel L. McCartney, Shane T. O’Sullivan, Karen Burr, Helen S. Torrance, Jonathan Phillips, Marion Bonneau, Susan M. Anderson, Paul Perry, Matthew Pearson, Costas Constantinides, Hazel Davidson-Smith, Mostafa Kabiri, Barbara Duff, Mandy Johnstone, H. Greg Polites, Stephen Lawrie, Douglas Blackwood, Colin A. Semple, Kathryn L. Evans, Michel Didier, Siddharthan Chandran, Andrew M. McIntosh, David J. Price, Miles D. Houslay, David J. Porteous, J. Kirsty Millar

## Abstract

The neuromodulatory gene *DISC1* is disrupted by a t(1;11) translocation that is highly penetrant for schizophrenia and affective disorders, but how this translocation affects DISC1 function is incompletely understood. N-Methyl-D-Aspartate receptors (NMDAR) play a central role in synaptic plasticity and cognition, and are implicated in the pathophysiology of schizophrenia through genetic and functional studies. We show that the NMDAR subunit GluN2B complexes with DISC1-associated trafficking factor TRAK1, while DISC1 interacts with the GluN1 subunit and regulates dendritic NMDAR motility in cultured mouse neurons. Moreover, in the first mutant mouse that models DISC1 disruption by the translocation, the pool of NMDAR transport vesicles and surface/synaptic NMDAR expression are increased. Since NMDAR cell surface/synaptic expression is tightly regulated to ensure correct function, these changes in the mutant mouse are likely to affect NMDAR signalling and synaptic plasticity. Consistent with these observations, RNASeq analysis of translocation carrier-derived human neurons indicates abnormalities of excitatory synapses and vesicle dynamics. RNASeq analysis of the human neurons also identifies many differentially expressed genes previously highlighted as putative schizophrenia and/or depression risk factors through large-scale genome-wide association and copy number variant studies, indicating that the translocation triggers common disease pathways that are shared with unrelated psychiatric patients. Altogether our findings suggest that translocation-induced disease mechanisms are likely to be relevant to mental illness in general, and that such disease mechanisms include altered NMDAR dynamics and excitatory synapse function. This could contribute to the cognitive disorders displayed by translocation carriers.

## INTRODUCTION

NMDAR are vital for synaptic plasticity and cognitive processes, and are strongly implicated in the pathophysiology of schizophrenia in particular^1^. *GRIN2A*, encoding the NMDAR GluN2A subunit was recently reported to show genome-wide significant association with schizophrenia^2^, while genomic copy number variants (CNVs) enriched in schizophrenia patients target *GRIN1*, encoding the GluN1 subunit^3, 4^. Such findings support a role for NMDAR in psychiatric disorders but direct mechanistic insight is still required.

NMDAR function is regulated at many stages including subunit expression and composition, and dynamic modulation of surface and synaptic levels. The latter can be elicited through control of NMDAR forward trafficking to the plasma membrane, subsequent insertion into synapses, or endocytosis^5^. The obligatory GluN1 subunit is incorporated into all NMDAR, along with other subunit types including GluN2A/B. GluN1 is synthesised in excess and stored within the endoplasmic reticulum (ER) where NMDAR are assembled prior to transportation to the Golgi and onwards to the cell surface^5^. Every stage of NMDAR forward trafficking is tightly modulated to ensure that the required quantity of receptors is present at the cell surface and synapse^5^. Genetic dysregulation of forward trafficking would be predicted to adversely affect NMDAR function, with knock-on effects for synapse strength and plasticity^5^.

DISC1 is disrupted in a large, multi-generation family by a chromosomal t(1;11) translocation that is linked to major mental illness^6–8^. DISC1 is critical for several processes in the developing and adult brain^9,10^, and has been connected to NMDAR function through regulation of downstream processes that underlie synaptic plasticity^11, 12^. DISC1 also regulates neuronal microtubule-based cargo transport, including trafficking of mitochondria, synaptic vesicles, and messenger RNAs^13–15^, and associates with the motor protein adaptors TRAK1 and TRAK2^16, 17^.

Here we demonstrate direct interaction between GluN1 and DISC1, and association between GluN2B and TRAK1. Applying a novel live-imaging method, we demonstrate that DISC1 regulates dendritic NMDAR motility. Utilising a mouse model of DISC1 disruption by the translocation we show that mutant mouse neurons exhibit increased NMDAR fast active transport and cell surface/synaptic NMDAR expression. These observations implicate dysregulated NMDAR dynamics/ signalling and excitatory synapse dysfunction in the psychiatric disorders displayed by translocation carriers. In further support of a general disease mechanism revealed by the translocation, we find that it impacts biological pathways highlighted by independent schizophrenia and depression GWAS and CNV studies.

## MATERIALS AND METHODS

Detailed materials and methods for all experiments are provided in the supplementary information. Essential information is included below.

### Generation of induced pluripotent stem cells (IPSC) from dermal fibroblasts

Dermal fibroblasts were reprogrammed using non-integrating episomal plasmids incorporating Oct3/4, shRNA to p53, SOX2, KLF4, L-MYC and LIN28^18^, and converted to neural precursor cells by dual-SMAD signaling inhibition^19^.

### Generation of a mouse model of the t(1;11) translocation

Mice were genetically engineered using Regeneron’s GEMM platform (VelociMouse^®^).*Disc1* was targeted in embryonic stem cell clones using a vector containing mouse *Disc1* genomic DNA encompassing exons 6, 7 and 8 as the 5’ homologous arm, and sequences 3’ of *Disc1* as the 3’ homologous arm. Human chromosome 11 genomic DNA, encompassing putative exons 4, 5, 6, 7a, and 7b of *DISC1FP1*^20^ followed by a loxP-flanked neo cassette, was inserted between the mouse homologous arms. Homologous recombination removed 98,550bp of mouse *Disc1* downstream of exon 8, and inserted 114,843bp of human *DISC1FP1* sequence. The edited endogenous mouse *Disc1* locus mimics the *DISC1/DISC1FP1* gene fusion event on the derived chromosome 1 in translocation carriers, and the mutation is referred to as *Der1*.

### RNA sequencing

Three control and three translocation-carrying neural precursor lines were each differentiated to neurons independently in triplicate. RNA was extracted and sequenced to a depth of 60 million paired-end reads. Differential gene expression was analysed at the whole gene and single exon levels using DESeq2 from the R statistical package^21^, and DEXSeq^22^, respectively.

### Dendra2 NMDAR trafficking assay

A Dendra2 tag was fused to the GluN1 C-terminus. Dendra2 exhibits green fluorescence, but can be photoconverted to red fluorescence. DIV7 hippocampal neurons were co-transfected with GluN1-Dendra2 plus HA-GluN2B to maximise NMDAR assembly, and therefore trafficking outwith the ER. Dendra2 was photoconverted within a region of interest positioned on a primary dendrite. Time-lapse images were captured every 15 seconds for three minutes and red fluorescence bidirectional movement along dendrites was quantified. Essentially, dendrites were divided into 5μm bins, and red fluorescence intensity was quantified within each bin at each timepoint. Finally, fluorescence intensity within each bin was normalised to dendritic area and fluorescence intensity in bin zero at time zero.

Fluorescence peak average velocities were determined individually for each neuron using the 10μm and 15μm bins where the peak was clearly identifiable. Maximum velocity estimates were determined individually for each neuron using the 25-40μm bins where the appearance of the leading edge of the wave of red fluorescence could be ascertained.

## RESULTS

### Neural precursor cells (NPC) and neurons derived from t(1;11) translocation carriers

Dermal fibroblasts from translocation carriers diagnosed with schizophrenia, recurrent major depression, cyclothymia or recurrent major depression plus bipolar disorder (not otherwise specified, and karyotypically normal family members (Supplementary Figure 1) were reprogrammed using non-integrating episomes^18^. NPC lines were generated from the IPSC (Supplementary Figure 2a-c) as described previously^19^. All NPC lines exhibit typical morphology and form rosettes in culture, with almost 100% of cells expressing the neural progenitor marker NES (Supplementary Figure 2c). Each NPC line consistently differentiates to produce neurons expressing the neuronal marker βIII-tubulin (Supplementary Figure 2d).

To examine the effect of the translocation upon DISC1 transcript levels in the t(1;11) family NPCs and neurons, NPCs were differentiated over five weeks, with sampling at time zero (before differentiation onset) and weekly thereafter, to obtain RNA for quantitative RT-PCR analysis. Wild-type DISC1 expression is decreased in translocation-carrying NPCs and neurons relative to controls (Figure 1a). Moreover, DISC1 expression increases as differentiation progresses (Figure 1a,b), although the proportional change each week is not altered by the translocation (Figure 1b). DISC1 expression may therefore be particularly important in neurons. An antibody that detects full-length DISC1 (Supplementary Figure 3a) confirmed its reduced expression in NPCs and neurons (Figure 1c,d).

**Figure 1.**
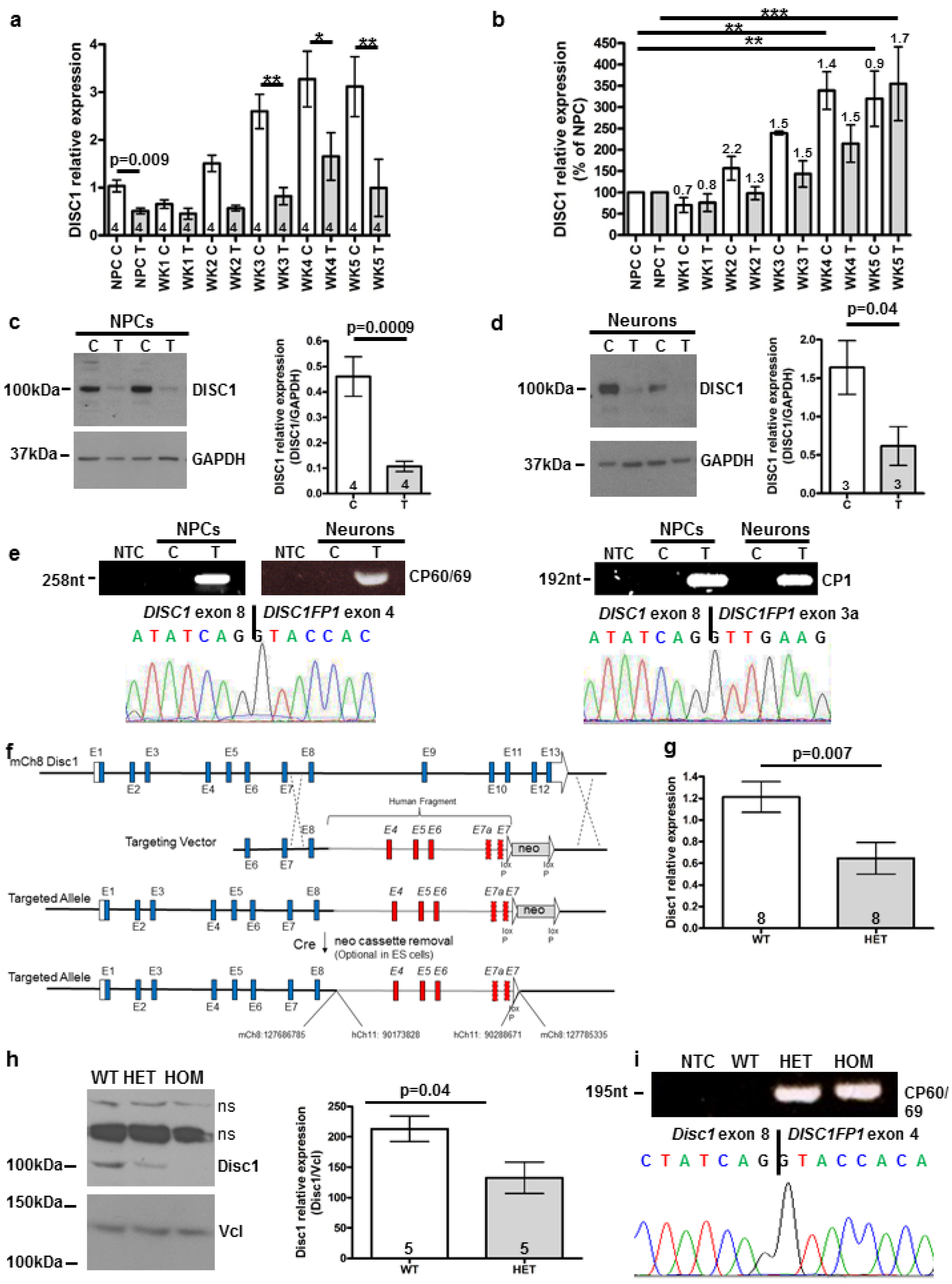
DISC1 expression in neural precursors and neurons derived from members of the t(1;11) family by the IPSC route, and in the *Der1* mouse model. (**a**) Quantitative RT-PCR analysis of DISC1 transcripts in NPCs and differentiating neurons over five weeks. Data analysed by two-tailed t-test (NPC samples only) and one-way ANOVA (all samples, p<0.0001), with pairwise Bonferroni post-test. (**b**) Data from a expressed as percentage of control. Data analysed by one-way ANOVA (p<0.0001) with pairwise Bonferroni post-test between NPC and each week of differentiation. Numbers above the columns indicate fold-change in comparison to the previous week. (**c**)(**d**) Immunoblotting of DISC1 protein in neural precursors, c, or NPCs differentiated to neurons in triplicate, d. Left, representative immunoblots; Right, densitometric analysis. Data analysed by two-tailed t-test. (**e**) Detection of chimeric CP60/CP69 and CP1 transcripts by RT-PCR and sequencing. (**f**) Schematic of the mouse endogenous *Disc1* allele targeting strategy. (**g**) Quantitative RT-PCR analysis of *Disc1* transcripts in adult mouse whole brain. Data analysed by two-tailed t-test. (**h**) Disc1 protein expression in adult mouse whole brain. Left, representative immunoblot; Right, quantification. Data analysed by two-tailed t-test. ns, non-specific bands (**i**) Detection of chimeric CP60/CP69 transcripts by RT-PCR and sequencing. NTC, non-template control; C, control; T, translocation carrier; WT, *Disc1*^*wt/wt*^; HET, *Disc1*^*wt/Der1*^; HOM, *Disc1*^*Der1/Der1*^; error bars represent SEM; * p<0.05; ** p<0.01; ***p<0.001; n indicated on graphs

Chimeric CP1 and CP60/69 transcripts, resulting from translocation-induced *DISC1* gene fusion to the *DISC1FP1* (otherwise known as Boymaw) gene on the derived chromosome 1^20^ were PCR-amplified from NPCs and neurons derived from translocation carriers (Figure 1e). These transcripts encode aberrant forms of DISC1 which, when exogenously expressed, are deleterious^20^, thus they are likely damaging to the brain *in vivo*. However, their endogenous expression awaits confirmation.

*DISC2* and *DISC1FP1*, also disrupted by the translocation, are expressed in the NPCs and neurons, but at low, non-quantifiable levels (Supplementary Figure 3b).

### Transcriptomic analysis of t(1;11) family neurons

We next performed RNA sequencing (RNASeq) analysis as an unbiased screen for the effects of the translocation upon gene expression in the human cortical neurons. The translocation lines used were derived from patients diagnosed with schizophrenia, recurrent major depression, or cyclothymia. The use of cross-diagnostic samples aimed to ensure that the major effects of the translocation are identified, rather than any disorder-specific effects due to genetic background. 1,256 (BaseMean>10) of 22,753 expressed genes were found to be differentially expressed (adjusted p<0.05, Figure 2a, Supplementary Table 1a). 938 genes also exhibit differential expression of individual exons (adjusted p<0.05), an indicator of altered isoform expression (Supplementary Table 1b).

**Figure 2.**
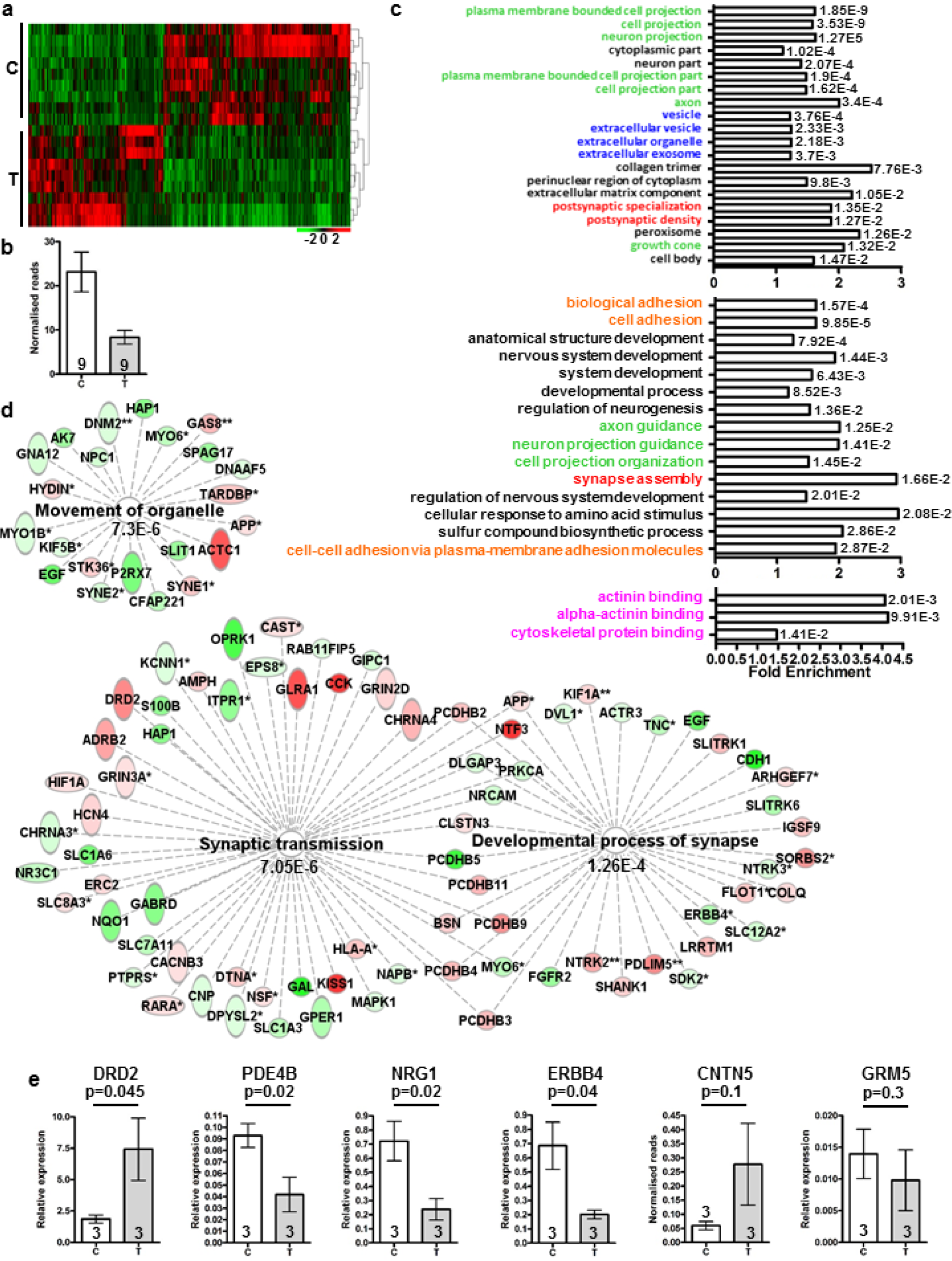
RNAseq analysis of human neurons. (**a**) Heat map of differentially expressed genes (DESeq2, adjusted p<0.05) in neurons from three control (28, 29, 30, Supplementary Figure 1) and three translocation lines (18, schizophrenia; 24, recurrent major depression; 55, cyclothymia), three differentiations each. (b) Normalised RNASeq reads spanning DISC1 exons 8 and 9 in all samples used for RNASeq analysis. (**c**) Top 20 significantly enriched GO Component (top), top 15 GO Process (middle), plus all GO Function (bottom) terms for combined DESeq2 plus DEXSeq lists, indicating FDR-corrected q values. Colours indicate related GO terms (**d**) Gene networks relating to synapses and organelle movement derived using DESeq2 plus DEXSeq lists in IPA. *genes identified by DEXSeq; ** genes identified by DEXSeq plus DESeq2; red, upregulated; green, downregulated (**e**) Quantitative RT-PCR confirms differential expression of *DRD2*, *PDE4B*, *NRG1* and *ERBB4*, but not of *CNTN5* and *GRM5*, using three control and three translocation lines differentiated to neurons in triplicate. Data analysed by one-tailed t-test. C, control; T, translocation carrier; error bars represent SEM; n indicated on graphs

IPSC-derived neuron cultures are a mix of cell types, thus it is possible that the translocation could affect gene expression patterns via influences upon cell fate. However analysis of genes whose expression was determined to be enriched in mouse or human astrocytes and neurons by single cell RNASeq analysis^20, 21^ found that only 7/41 mouse and 3/21 human astrocyte markers, and 8/25 mouse and 5/34 human neuron markers are dysregulated (Supplementary Table 1c,d). Four of the dysregulated mouse neuron markers, DLX1, DLX2, DLX5 and DLX6, are already known to be regulated by DISC1, indicating a subtle shift in neuronal properties^22,23^. Classic markers of neurons, and of inhibitory GABAergic neurons, are also unaffected (Supplementary Table 1d). Moreover, analysis of genes whose expression is enriched in specific ‘communities’ of human excitatory and inhibitory neurons^21^ found little indication of change to either type overall (Supplementary Table 1e). Thus, the translocation does not affect the proportion of astrocytes versus neurons, or of excitatory versus inhibitory neurons, although neuron properties may be subtly altered. This is consistent with a previous study which examined the effects of a DISC1 mutation upon gene expression in IPSC-derived neurons^22^.

300 and 71 of the dysregulated genes, respectively, are also differentially expressed in IPSC-derived neurons carrying a frameshift mutation in *DISC1* exon 12^23^ (hypergeometric probability p=2.5E-21 Supplementary Table 1a,b) or exon 8^22^ (p=0.00002), indicating that independent *DISC1* mutations have overlapping effects. Differential *DISC1* expression was not initially detected in the whole gene analysis, presumably due to the presence of chimeric *DISC1* transcripts originating from both derived chromosomes (Figure 1e, Supplementary Figure 3c). However comparison of reads that span intron 8, the location of the translocation breakpoint, and that can therefore only arise from wild-type transcripts, indicates that wild-type *DISC1* transcripts are indeed reduced (Figure 2b).

Due to the potential for positional, or other, effects of the translocation upon neighbouring gene expression, we examined transcript levels over a region of approximately 20Mb proximal and distal to the translocation breakpoints on chromosomes 1 and 11. Such analysis identified a number of differentially expressed genes and exons, and possible clustering of dysregulated genes around the chromosome 11 breakpoint (Supplementary Table 1f, g).

Gene ontology (GO) analysis (Figure 2c) highlighted cell/neuron projections (7 of the top 20 GO Component terms), the post-synaptic density of excitatory synapses (2/20) and vesicles/organelles (4/20). Additional significant GO Process and GO Function terms relate to synapse assembly, cell adhesion and the actin cytoskeleton. Ingenuity Pathway Analysis (IPA), a complementary method, identified similar terms, including terms that predict the translocation will affect synapse assembly and function in mature neurons of the brain (Figure 2d). IPA also found the term ‘movement of organelle’ was significant, (Figure 2d). Given that the RNAseq analysis is unbiased, it is notable that *DISC1* regulates processes related to three of the major enriched categories; neurite extension^10^; synaptic vesicle and mitochondrial trafficking^13,14^; and actin remodelling^11^. We therefore infer that these process-related gene expression changes are most likely due to DISC1 disruption.

While the translocation is unique to a single family, translocation carriers do not present unusual clinical symptoms outside current diagnostic criteria, thus the consequences of the translocation may converge upon biological processes shared with other unrelated patients. In support of this, several of the differentially expressed genes in t(1;11) neurons correspond to genome-wide association study (GWAS) findings for schizophrenia^2,24^. Expression of 33 of the 348 genes from 25 the 108 associated loci (hypergeometric probability p=7E-6 indicates enrichment for loci containing at least one dysregulated gene) identified in the first study, and of 37 of the 481 genes from 24 of the 145 loci (hypergeometric probability p=0.001) identified in the second study, is altered at the whole gene and/or exon level in control versus translocation lines (Supplementary Table 1a,b). Of these dysregulated putative schizophrenia genes, changed expression of *DRD2*, encoding the dopamine D2 receptor, a target of all antipsychotics in clinical use, and *PDE4B*, encoding a cAMP-degrading phosphodiesterase and known DISC1 interactor^25^ was confirmed by quantitative RT-PCR (Figure 2e). Altered expression was also confirmed for genes encoding the historical candidate *NRG1*^26^ and its receptor *ERBB4* (Figure 2e), both of which have previously been linked functionally to DISC1^27,28^. Expression of 12 of 52 genes encoding synaptic proteins that are targeted by recurrent CNVs in schizophrenia^3^ is also altered (hypergeometric probability p=0.001, Supplementary Table 1a,b). Four of 70 genes at four of 44 depression-associated loci (9.1% of loci, no enrichment)^29^ identified through GWAS are also dysregulated (Supplementary Table 1a,b).

Finally, three of four recently identified putative modifier loci in the t(1;11) family^30^ are adjacent to the translocation breakpoints on chromosome 1 or 11 (Supplementary Table 1f,g). Of the three genes of interest identified therein, *GRM5* is not dysregulated in the human cortical neurons (Figure 2e, Supplementary Table 1a,b), *CAPN8* is expressed at very low, non-quantifiable levels, and *CNTN5* is dysregulated according to the RNASeq analysis (p=0.0008, Supplementary Table 1a,b). However, although quantitative RT-PCR also suggests increased *CNTN5* expression, this did not achieve statistical significance (Figure 2e). The fourth locus, encompassing *PDE4D*, could not be examined because the cell lines used for RNASeq analysis and quantitative RT-PCR do not carry the required haplotype.

### A mutant mouse recapitulates the effect of the t(1;11) translocation upon *DISC1*

A mutant mouse was generated by deleting genomic DNA containing *Disc1* exons 9-13, and replacing it with human chromosome 11 genomic DNA containing relevant *DISC1FP1* exons (Figure 1f). This mimics the gene fusion caused by the translocation on the derived chromosome 1^20^, thus we refer to the mutation as *Der1*.

Quantitative RT-PCR demonstrated that wild-type *Disc1* transcript expression is reduced by approximately half in whole brain isolated from adult heterozygous *Der1*(*Disc1*^*wt/Der1*^) mice in comparison to wild-type (*Disc1*^*wt/wt*^) mice (Figure 1g). Full-length *Disc1* protein expression is detected at approximately 100kDa, and is correspondingly reduced in *Disc1*^*wt/Der1*^ versus *Disc1*^*wt/wt*^ mice (Figure 1h). As expected wild-type *Disc1* transcripts and protein were undetectable in whole brain from homozygous (*Disc1*^*Der1/Der1*^) mice (data not shown, Figure 1h, respectively). CP60/69 transcripts were detected using RT-PCR in whole brain from adult *Disc1*^*wt/Der1*^ and *Disc1*^*Der1/Der1*^ mice (Figure 1i), and quantitative RT-PCR in *Disc1*^*Der1/Der1*^ mice demonstrated widespread expression of these transcripts (Supplementary Figure 3d). Quantitative analysis of wild-type *Disc1* expression in *Disc1*^*wt/wt*^ mice generated a similar pattern of expression to that of the chimeric transcripts (Supplementary Figure 3d), indicating that *Disc1* promoter activity is not greatly affected by the *Der1* mutation. Further characterisation of the *Der1* mutant mouse will be published elsewhere.

### Interaction between DISC1 and GluN1

Immunoprecipitation experiments using endogenous brain proteins have previously demonstrated that GluN1 and Disc1 exist in shared complexes *in vivo*^31^. Here, using FLAG-DISC1 and HA-tagged GluN1, we confirmed that they can be co-immunoprecipitated using COS7 cells (Figure 3a). We next used FLAG-DISC1 to probe peptide arrays of the GluN1 cytoplasmic tail, encompassing the alternatively spliced C0, C1 and C2 cassettes (Figure 3b)^32^. FLAG-DISC1 was found to bind directly to GluN1 peptide spots 1-16, encompassing amino acids 831-930 and, most strongly, to three regions, namely within peptides 6-8, 13 and 16, corresponding to the C1, C2 and C0 cassettes. To confirm DISC1 binding to these regions we demonstrated that FLAG-DISC1 efficiently co-immunoprecipitates GST-tagged GluN1 C0-C1-C2 from COS7 cell lysates (Figure 3c). All known GluN1 isoforms contain at least one of these cassettes^32^ and thus can potentially interact with DISC1.

**Figure 3.**
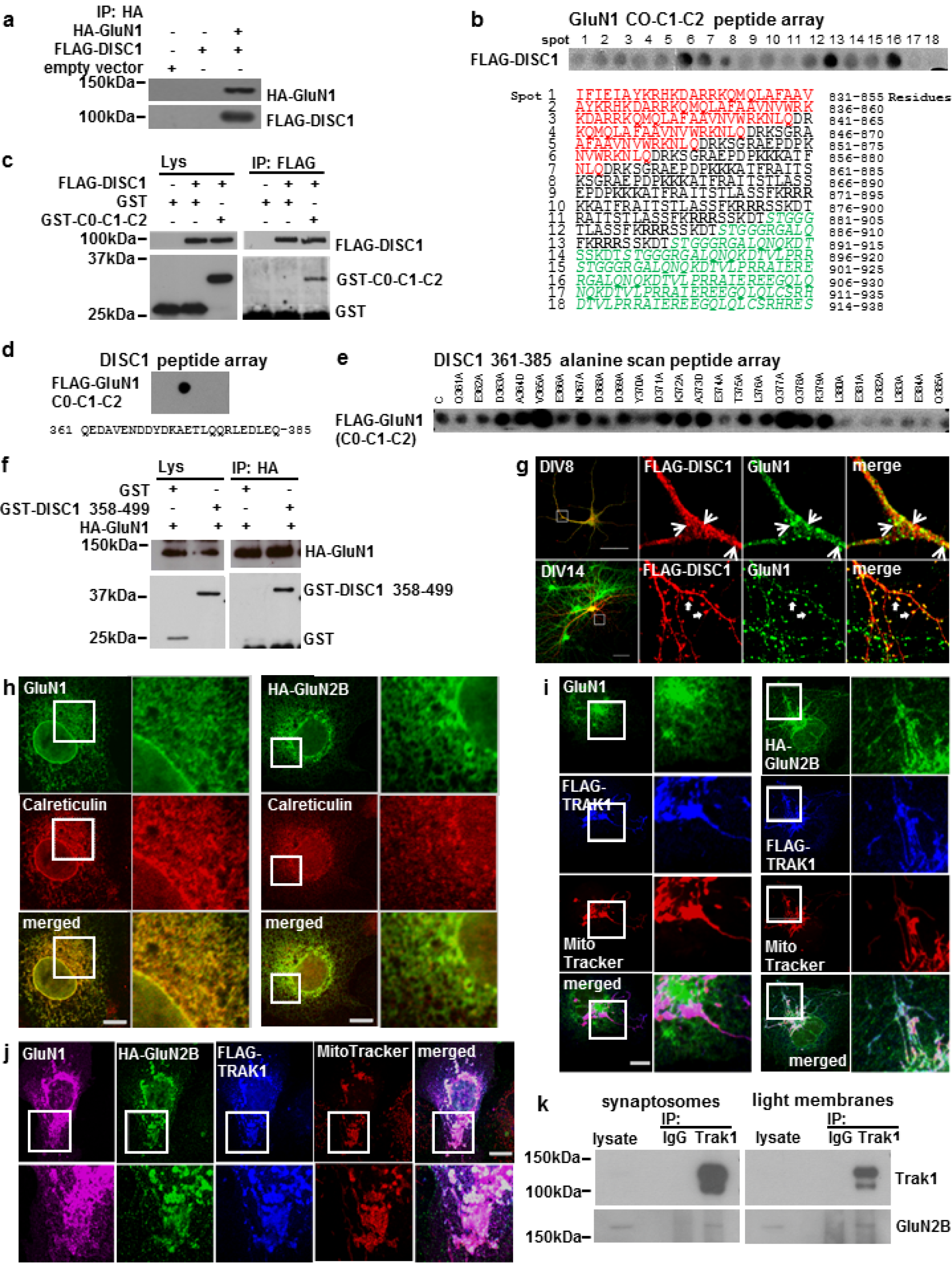
DISC1 interacts with GluN1, TRAK1 associates with GluN2B. (**a**) HA-GluN1 co-immunoprecipitates FLAG-DISC1 from transfected COS7 cells. (**b**) Upper, GluN1 cytoplasmic tail (C0-C1-C2) peptide array hybridised with Flag-DISC1. Each spot represents a single peptide. red, black and green text marks the C0, C1 and C2 cassettes respectively. ER retention signals are in bold. (**c**) FLAG-DISC1 co-immunoprecipitates GST-tagged C0-C1-C2. (**d**) DISC1 peptide array probed with FLAG-C0-C1-C2. (**e**) DISC1 361-385 GluN1 binding region alanine scan probed with FLAG-C0-C1-C2. C, unmutated peptide. (**f**) HA-GluN1 co-immunoprecipitates GST-DISC1 amino acids 358-499 from transfected COS7 cells. (**g**) Co-localisation of FLAG-DISC1 and endogenous GluN1 in cultured DIV8 and DIV14 hippocampal neurons. Arrowheads and arrows indicate example sites of colocalisation. (**h**) COS7 cells transfected with GluN1 or HA-GluN2B expression constructs were labelled with antibodies specific for GluN1 or HA, plus the ER marker Calreticulin. (**i**) COS7 cells co-transfected with FLAG-TRAK1 plus GluN1 (left) or HA-GluN2B (right) expression constructs were labelled using antibodies specific for GluN1 or HA, plus antiFLAG and the mitochondrial dye Mitotracker CMXRos. (**j**) COS7 cells triple-transfected with FLAG-TRAK1, GluN1 plus HA-GluN2B expression constructs were labelled using antibodies specific for GluN1, HA and FLAG, plus Mitotracker CMXRos. COS7 cells were used because they are ideal for exogenous protein expression due to their high transfection efficiency, large size and low profile, which facilitate co-immunoprecipitation and colocalisation studies to complement endogenous protein studies in neurons. (**k**) Trak1 immunoprecipitates GluN2B from adult mouse brain synaptosome and light membrane fractions. Scale bars, 50μm in G, otherwise 20μm; white boxes indicate enlarged areas

Reciprocal probing of a human DISC1 peptide array with the cytoplasmic tail of GluN1 fused to FLAG revealed a GluN1 contact site between amino acids 361-385 (Figure 3d). To confirm this, an alanine scanning peptide array of DISC1 amino acids 361-385 was interrogated with the same GluN1 probe (Figure 3e), revealing that binding to the wild-type sequence is substantially weakened by alanine conversion of any one of amino acids E374, L380, E381, D382, L383, E384 and Q385. The DISC1 head domain therefore contacts GluN1 via a single site in which residues 380-385 are critical for binding. This binding site was independently confirmed by co-immunoprecipitating HA-GluN1 and GST-DISC1 amino acids 358-499 from COS7 cells (Figure 3f). Moreover, exogenous DISC1 co-localises with endogenous GluN1 in cultured DIV8 and DIV14 hippocampal neurons (Figure 3g). These observations indicate that DISC1 and the putative schizophrenia risk factor GluN1^3^ have the capacity to interact directly *in vivo*.

### TRAK1 complexes with GluN2B

Because DISC1 interacts with GluN1 and regulates cargo transport, we speculated that DISC1 and TRAK1 might be involved in NMDAR trafficking. To test this, we co-transfected COS7 cells with TRAK1 plus NMDAR subunit GluN1 or GluN2B expression constructs. In the absence of TRAK1, both GluN1 and HA-GluN2B predominantly co-localise with the ER marker Calreticulin (Figure 3h), while FLAG-TRAK1 is strongly targeted to mitochondria, as expected^17^ (Figure 3i). In co-transfected cells there is no appreciable co-localisation between FLAG-TRAK1 and GluN1 (Figure 3i), but in the presence of FLAG-TRAK1, HA-GluN2B redistributes from the ER to mitochondria (Figure 3i). Moreover, in triple transfected cells, in the presence of HA-GluN2B plus FLAG-TRAK1, a substantial proportion of GluN1 localises to mitochondria (Figure 3j), indicating that assembled NMDAR can associate with TRAK1. We next immunoprecipitated Trak1 from mouse brain synaptosome and light membrane fractions where both proteins are enriched^33^ (data not shown), and demonstrated co-precipitation of GluN2B (Figure 3k, Supplementary Figure 4). The DISC1-associated trafficking factor Trak1 thus associates with the NMDAR GluN2B subunit *in vivo*.

DISC1, with TRAK1, is now well established as a regulator of neuronal intracellular trafficking^34^. The GluN1/ DISC1/TRAK1 and TRAK1/GluN2B associations therefore imply that DISC1 regulates motility of GluN2B-containing NMDAR.

DISC1 dysregulates dendritic GluN1-Dendra2 motility in mouse hippocampal neurons To examine NMDAR motility in neurons we fused the green fluorescent protein Dendra2 to GluN1, which is incorporated into all NMDAR. Dendritic GluN1-Dendra2 has a granular appearance similar to that of endogenous GluN1 (Figure 4a, Supplementary Figure 5a)^35^. Dendra2 can be stably photoconverted to red fluorescence, enabling tracking of a specific red GluN1 population (Supplementary Video). Dendra2 fusion to the GluN1 C-terminus does not interfere with receptor assembly or trafficking to synapses because GluN1-Dendra2 co-localises with the post-synaptic density marker PSD95 within dendritic spines (Supplementary Figure 5b). This can only occur following proper receptor assembly^5^.

**Figure 4.**
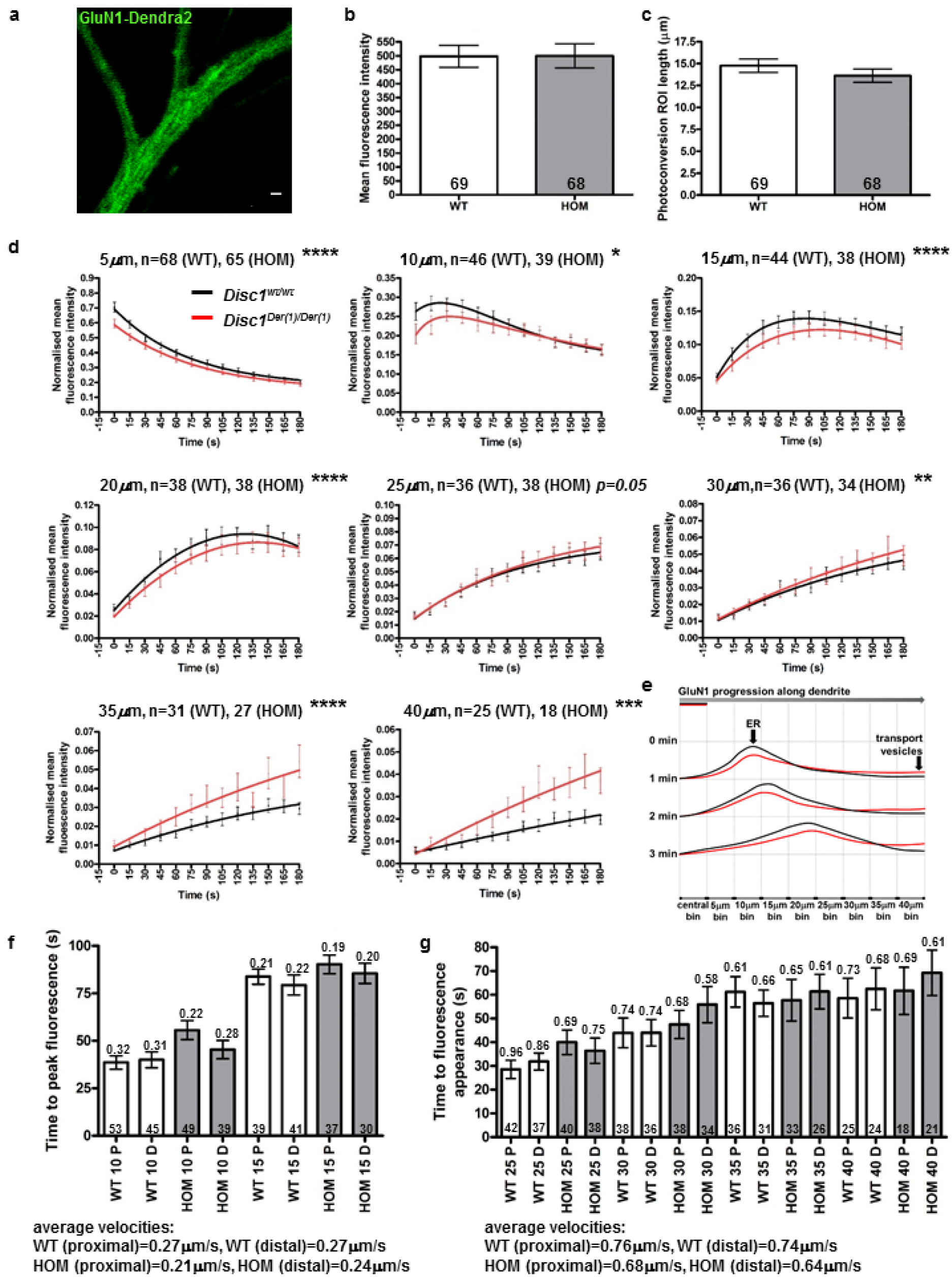
Altered distal dendritic NMDAR trafficking in *Disc1*^*Der1/Der1*^ hippocampal neurons. (**a**) Green Dendra2 fluorescence in dendrites of a DIV8 mouse hippocampal neuron transfected with GluN1-Dendra2 plus HA-GluN2B. Scale bar, 5μm (**b**)(**c**) Mean red fluorescence intensity, b, or ROI dendritic length, c, in the central bin in *Disc1*^*wt/wt*^ and *Disc1*^*Der1/Der1*^ DIV8 neurons was equal at time zero following photoconversion. (**d**) Quantification of fluorescence intensity over time in successive 5μm dendritic bins distal to the centre of the photoconversion ROI. Data analysed by timepoint-paired two tailed t-test. (**e**) Model of dendritic GluN1-Dendra2 motility. Photoconverted GluN1-Dendra2 progresses in a wave-like fashion, with the fastest and slowest moving GluN1-Dendra2 at the leading and trailing edges, respectively, and the bulk travelling as the ‘crest’. (**f**) Fluorescence peak velocity estimates for the 10μm and 15μm bins. Average time to peak fluorescence was converted to velocity, indicated above each bar. Average velocities were determined from the two bins. (**g**) Fast-moving GluN1-Dendra2 maximum velocity estimates for the 25μm-40μm bins. Average time to fluorescence appearance was converted to velocity, indicated above each bar. Average velocities were determined from the four bins. WT, *Disc1*^*wt/wt*^; HOM, *Disc1*^*Der1/Der1*^; error bars represent SEM; **** p<0.0001; *** p<0.001; ** p<0.01; *p<0.05; n indicated on graphs

Bulk GluN1-Dendra2 movement was quantified in cultured wild-type hippocampal neurons expressing exogenous human DISC1, at Days In Vitro 8 (DIV8), an age at which the dendritic arbour is suitably developed, but not too complex. Following GluN1-Dendra2 photoconversion, red fluorescence bidirectional spreading was imaged and analysed every 15 seconds for a total of 2 minutes in successive 5μm bins along the dendrites (Supplementary Figure 5c-n).

We examined the effect of wild-type DISC1 and of a change of arginine to tryptophan at position 37 (37W) in DISC1 (Supplementary Figures 6a-f,7a,7b). This sequence variant has been reported in four individuals, all with a diagnosis of psychiatric illness^36,37^, and dysregulates DISC1/TRAK1 association^17^. Wild-type DISC1 overexpression increases distal-moving red fluorescence in the 20-35μm bins (Supplementary Figure 6c,d), while in the proximal direction it reduces red fluorescence intensity in all bins (Supplementary Figure 7a,b). In contrast, DISC1-37W overexpression increases and decreases distal-moving red fluorescence in the 5-15μm and 25-35μm bins, respectively (Supplementary Figure 6c,d), while proximal-moving red fluorescence increases in the 5-20μm bins (Supplementary Figure 7a,b).

These data are consistent with the existence of at least two classes of distal-moving dendritic GluN1-Dendra2 in young hippocampal neurons that are differentially influenced by DISC1; 1) a slow-moving population in the 5-15/20μm bins (average velocity=0.2μm/s in empty vector-transfected neurons, Supplementary Figures 6d,6e,7b), representing the majority of the GluN1-Dendra2, and 2) a fast-moving population in the 25-35μm bins (average velocity=0.82μm/s in empty vector-transfected neurons, Supplementary Figures 6d,6f,7b).

These populations represent the trailing edge plus ‘crest’, and the leading edge, respectively, of a wave of motile GluN1-Dendra2 (Supplementary Figures 6d,7b). Our evidence for these populations aligns well with estimates of 0.2-0.3μm/s for movement of dendritic ER vesicles^38^ and with estimates of 0.76μm/s for the motility rate of assembled NMDAR-containing vesicles undergoing fast active transport in dendrites^39^, thus the slow and fast populations likely represent the ER GluN1 pool and NMDAR-containing vesicles undergoing active transport, respectively.

### Altered dendritic NMDAR motility in *Der1* mouse hippocampal neurons

We next quantified bulk GluN1-Dendra2 movement within neuronal dendrites from *Disc1*^*wVwt*^ and *Disc1*^*Der1/Der1*^ mice using the same method (Figure 4b-g, Supplementary Figure 7c,d). The same two pools of GluN1 were detected, with the distal-moving ER and transport vesicle pools respectively decreased and increased in *Disc1*^*Der1/Der1*^ mutant neurons (Figure 4d,e). A similar, but less marked, pattern of proximal GluN1-Dendra2 motility was observed (Supplementary Figure 7c,d).

#### Altered dendritic NMDAR surface distribution in *Der1* mouse hippocampal neurons

The NMDAR trafficking data indicate that DISC1 modulates NMDAR motility and consequently predict dysregulated cell surface NMDAR expression in neurons cultured from the *Der1* mutant mouse. To test this we quantified NMDAR cell surface expression in DIV21 cultured *Disc1*^*wt/wt*^, *Disc1*^*wt/Der1*^ and *Disc1*^*Der1/Der1*^ hippocampal neurons, an age at which neurons have matured and developed synapses. Cell surface expression of the NMDAR subunits GluN1, GluN2A and GluN2B, and of the post-synaptic density marker PSD95 and βIII-tubulin (Tuj1) was imaged at high resolution by three-dimensional Structured Illumination Microscopy (3D-SIM, Supplementary Figure 8) which achieved a lateral image resolution of 180nm for the NMDAR subunits and 120nm for PSD95.

3D reconstruction of βIII-tubulin and of GluN1, GluN2A and GluN2B was used to estimate dendritic volume, and the number and volume of individual surface puncta of each NMDAR subunit, respectively (Figure 5a,b, Supplementary Table 2). This analysis demonstrated increased density of GluN1 and GluN2B puncta in *Disc1*^*wt/Der1*^ and *Disc1*^*Der1/Der1*^ mutant neurons, while GluN2A puncta density is increased only in *Disc1*^*Der1/Der1*^ mutant neurons. Total surface expression is correspondingly increased in *Disc1*^*wt/Der1*^ and *Disc1*^*Der1/Der1*^ mutant neurons. Average puncta volumes are increased for all three NMDAR subunits, but only in *Disc1*^*wt/Der1*^ neurons, suggesting that NMDAR form larger clusters on the cell surface in heterozygous neurons. This observation supports the existence of putative aberrant proteins encoded by the chimeric CP60/69 transcripts. These proteins retain a region that is sufficient for self-association^40^, so there is potential for dominant-negative effects due to formation of multimers consisting of wild-type and mutant protein, a situation that occurs only in heterozygous cells.

**Figure 5.**
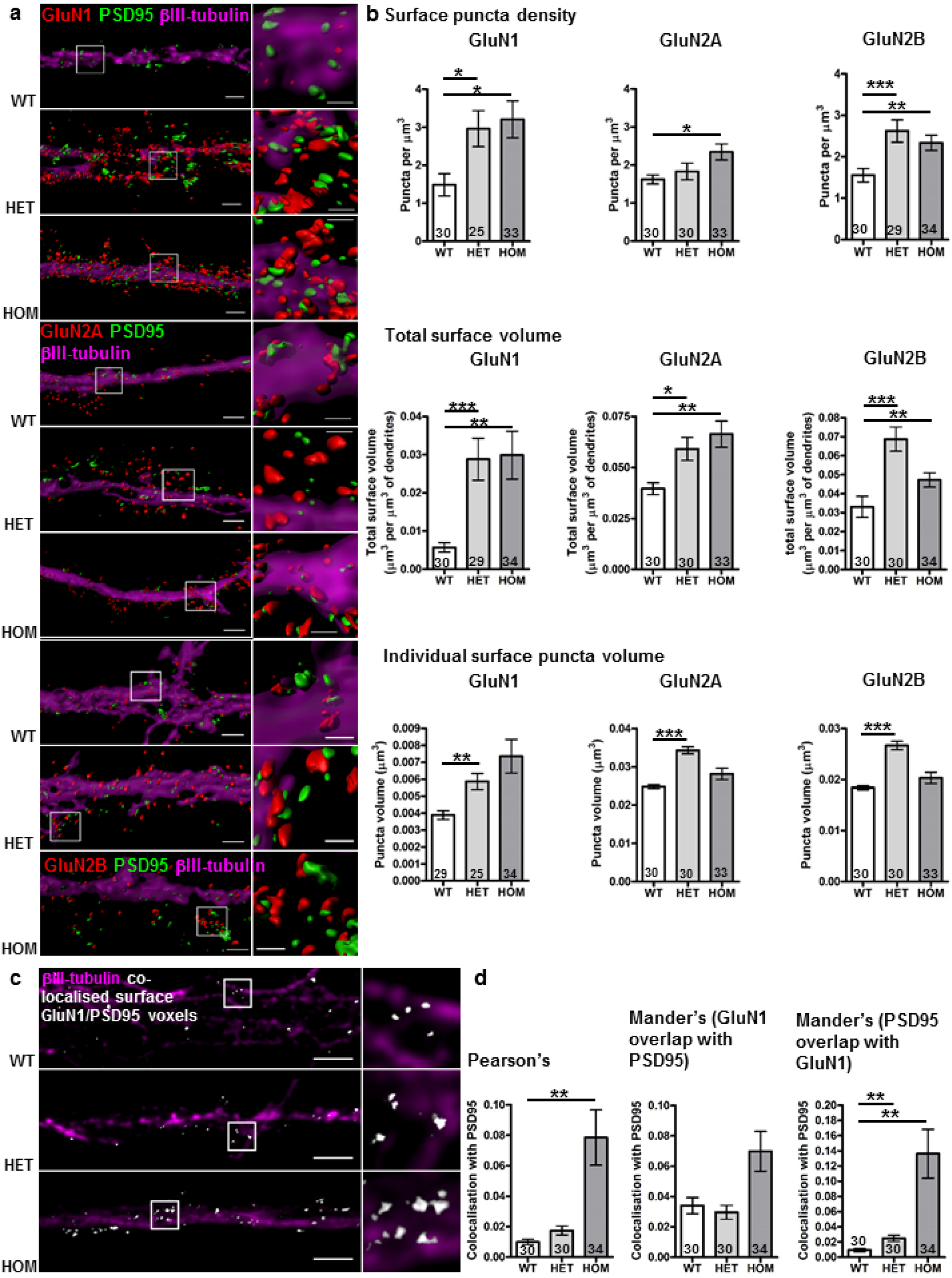
Altered dendritic NMDAR surface expression and GluN1 localisation to the post-synaptic density in *Disc1*^*wt/Der1*^ and *Disc1*^*Der1/Der1*^ hippocampal neurons. (**a**) Objects in 3D-SIM images visualised using the Imaris Isosurface tool. Touching objects are separated, with boundary lines between touching objects visible in the enlarged images. scale bars, 2μm in the full-size images, 0.6μm in enlarged insets indicated by white boxes (**b**) GluN1, GluN2A or GluN2B surface puncta density, total surface volume and individual surface puncta volume (all normalised to dendritic segment volume) from 3D reconstructions of primary dendrite segments of cultured DIV21 hippocampal neurons. Data analysed by Kruskal-Wallis (puncta density p=0.02, p=0.03, p=0.0004, total volume p=0.0025, p=0.4, p=0.003, puncta volume p=0.003, p<0.0001, p<0.0001 for GluN1, GluN2A and GluN2B respectively) followed by Dunn’s multiple comparison test. (**c**) Reconstructed 3D-SIM images of dendrites. Colocalised voxels, which contain signal from both PSD95 and surface-expressed GluN1 are shown in white. Scale bars, 3μm (**d**) GluN1 co-localisation with the PSD95 was evaluated on three measures. Pearson’s and Mander’s coefficients respectively indicate overall correlation of each signal or amount of GluN1 signal co-localised with PSD95 signal, and vice versa. Data analysed by Kruskal-Wallis (p=0.004, p=0.08, p=0.001 for Pearson’s, Mander’s M1 and Mander’s M2) followed by Dunn’s multiple comparison test. WT, *Disc1*^*wt/wt*^; HET, *Disc1*^*wt/Der1*^; HOM, *Disc1*^*Der1/Der1*^; error bars represent SEM; *** p<0.001; ** p<0.01; * p<0.05; n indicated on graphs

At the neuronal cell surface NMDAR are present at synaptic, perisynaptic and extrasynaptic sites^5^, thus only a proportion of receptors co-localise with PSD95 at synapses. Applying Pearson’s co-localisation coefficient, which quantifies the overall correlation between two fluorescent signals, indicates that PSD95 co-localisation with GluN1 is substantially increased in *Disc1*^*Der1/Der1*^ mutant neurons (Figure 5c,d). However, synaptic localisation of GluN2A and GluN2B is not greatly altered (Supplementary Figure 9a,b). Since GluN1 subunits cannot be transported to synapses unless they have assembled with other subunit types, we suggest that this discrepancy is related to the presence of two GluN1 subunits in every NMDAR, with variable inclusion of other subunit types, resulting in a clear effect for the former, but not for the latter. If correct, the GluN1/PSD95 co-localisation data indicate that NMDAR are more abundant at synapses in the *Disc1*^*Der1/Der1*^ mutant neurons.

Hippocampus protein extract immunoblotting demonstrated that total expression of GluN1, GluN2A and GluN2B is unaltered by the mutation in adult *Disc1*^*wt/Der1*^ or *Disc1*^*Der1/Der1*^ mice (Supplementary Figure 9c), confirming that surface expression changes are due to altered NMDA receptor subunit distribution.

#### Altered distribution of PSD95 in *Der1* hippocampal mouse neurons

PSD95 is one of the most abundant molecular scaffolds at the postsynaptic density of excitatory synapses. It has many functions including organisation of the postsynaptic density, NMDAR stabilisation at synapses, assembly of NMDAR-associated signalling complexes and recruitment of AMPA receptors (AMPAR), the major excitatory receptors in the brain^41^. PSD95 is thus a major determinant of excitatory synapse function and neurotransmission. Expression of PSD95 was therefore quantified in the imaged set of hippocampal neurons to determine whether synapse size and structure is also affected by the *Der1* mutation.

Super-resolution microscopy has demonstrated the existence of varying numbers of PSD95 puncta (nanodomains) at the post-synaptic density^42–44^. Most synapses have a single PSD95 nanodomain, but some have multiples^42–44^. Using 3D-SIM, PSD95 is visible as a combination of single spots and distinct clusters of multiple puncta (Figure 6a, Supplementary Figure 7), indicating that this super-resolution technique is able to resolve PSD95 nanodomains within clusters at the postsynaptic density. Total dendritic PSD95 volume is not affected by the *Der1* mutation (Figure 6b), however its distribution is altered. Analysis of the number of nanodomains per cluster (Supplementary Figure 8b-f) found that *Disc1*^*wt/Der1*^ and *Disc1*^*Der1/Der1*^ neurons have an increased density of single nanodomains (Figure 7c), and of small nanodomain clusters of 2-3 (Figure 7d).

**Figure 6.**
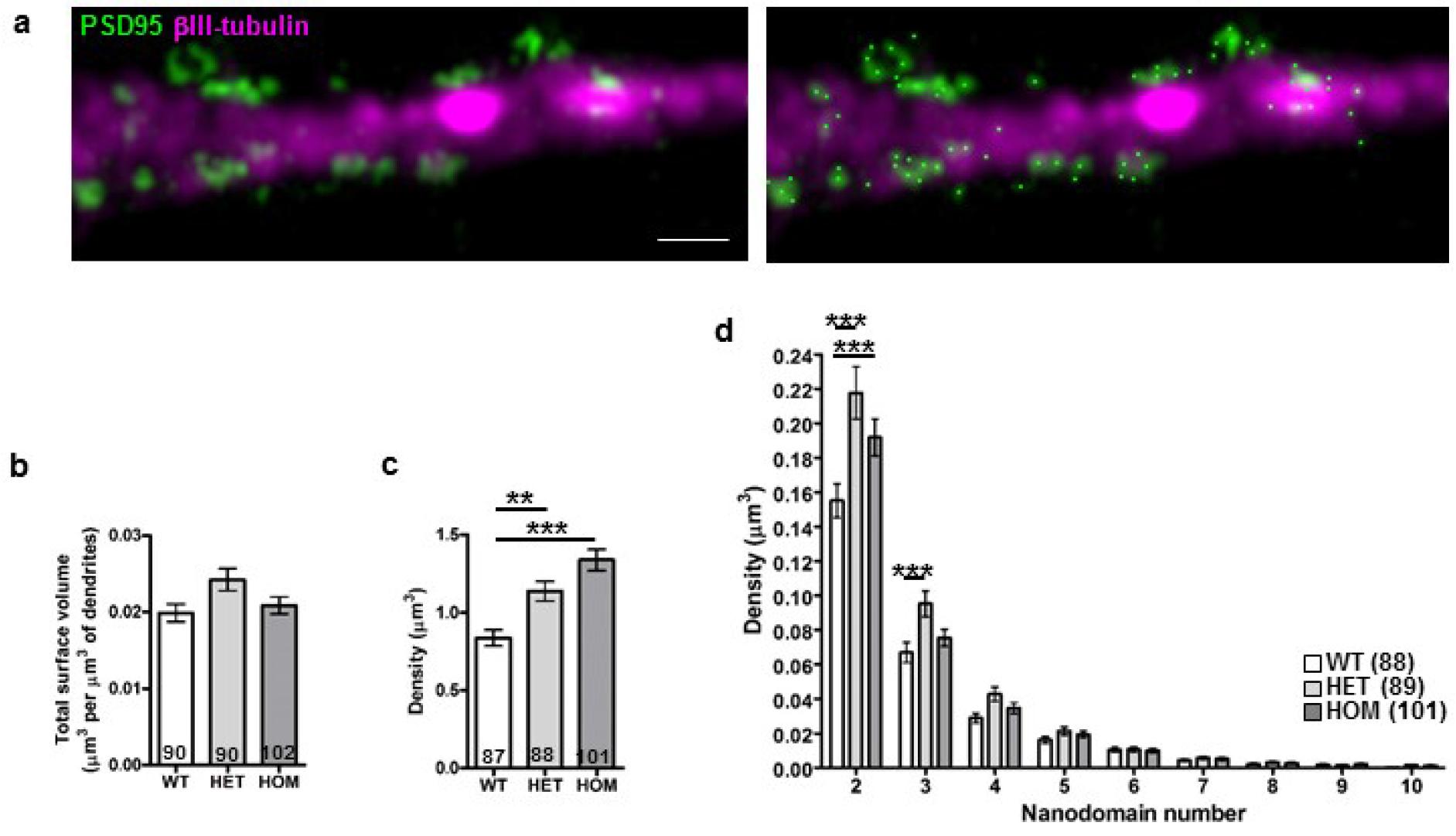
Altered PSD95 distribution in *Disc1*^*wt/Der1*^ and *Disc1*^*Der1/Der1*^ hippocampal neurons. (**a**) Left, raw 3D-SIM image of a dendrite with PSD95 clusters and nanodomains. Right, the same image with nanodomain centres (bright green spots) identified using Imaris. Scale bar, 1μm, (**b**) PSD95 total volume normalised to dendritic segment volume. Data analysed by Kruskal-Wallis (not significant). (**c**) Quantification of single PSD95 nanodomains. Data analysed by one-way ANOVA (p<0.0001) (**d**) Quantification of PSD95 nanodomain number per cluster. Data analysed by two-way ANOVA which found a significant interaction between genotype and nanodomain number per cluster (p<0.0001). WT, *Disc1*^*wt/wt*^; HET, *Disc1*^*wt/Der1*^; HOM, *Disc1*^*Der1/Der1*^; error bars represent SEM; *** p<0.001; ** p<0.01; n indicated on graphs

### DISCUSSION

Recent large-scale GWAS and CNV studies implicate excitatory synapses, NMDAR subunits GluN1 and GluN2A, and synaptic plasticity as causal factors in schizophrenia^2–4,24^. We show here that multiple genes identified in these studies are differentially expressed in human neurons carrying a t(1;11) linked to major mental illness. Moreover, the gene *DISC1*, disrupted by the t(1;11), encodes a direct interactor of the NMDAR obligatory GluN1 subunit. This convergence of the t(1;11) with schizophrenia risk genes indicates that the t(1;11) is a trigger for disease pathways which are shared with unrelated schizophrenia patients. Affective disorders are also prevalent in t(1;11) carriers and we found differential expression of putative depression risk factors in the t(1;11)-carrying human neurons, although not as strikingly as for schizophrenia.

We demonstrate that DISC1 regulates NMDAR motility within dendrites, a process that is critical for controlling cell surface and synaptic NMDAR expression. NMDAR motility is influenced by a DISC1 37W mutation^36, 37^, previously shown to affect mitochondrial motility^17^, and by the mouse *Der1* mutation which accurately models the effect of the t(1;11) upon DISC1 expression. DISC1, DISC1-37W and the *Der1* mutation influence the size of the GluN1 ER pool, indicating, in the absence of altered NMDAR subunit expression, that DISC1 regulates NMDAR ER-exit. Indeed DISC1 is ER-associated^16^, and may directly impact NMDAR ER-exit because the binding site for DISC1 on GluN1 encompasses two ER retention signals (Figure 3b)^5^. However, a simple model whereby increased/decreased ER-exit leads to respectively decreased/increased transport pool size cannot fully explain the effects of DISC1 upon NMDAR motility due to the lack of an overt relationship between the observed sizes of the ER and active transport pools. We therefore propose that DISC1 influences the active transport pool independently of ER-exit, via TRAK1, which associates with the NMDAR subunit GluN2B, and its role as a regulator of neuronal cargo trafficking^34^.

Mouse neurons expressing DISC1-37W exhibit a decreased pool of NMDAR undergoing fast active transport, while *Der1* mutant mouse neurons exhibit an increased pool. Since the 37W variant has to date been found only in psychiatric patients, this suggests that dysregulation of this pool, up or down, is deleterious. Upregulation of the pool in *Der1* mutant mouse neurons correlates strikingly with the augmented density and overall volume of cell surface NMDAR clusters, suggesting that increased trafficking/membrane insertion largely account for the greater surface NMDAR volume. NMDAR are constantly exchanged between synaptic and extrasynaptic sites^45^, with synaptic insertion events believed to result from stochastic postsynaptic density-mediated capture of receptors undergoing lateral diffusion within the cell membrane^46^. Consequently, the elevated NMDAR surface volume in *Der1* mutant mouse neurons likely increases the probability of their diffusion to, and insertion into, synapses, thus explaining the increased synaptic localisation of GluN1.

In addition to the NMDAR abnormalities, the post-synaptic density scaffold PSD95 is aberrantly distributed in *Der1* mutant mouse neurons. Postsynaptic density size at excitatory synapses is directly related to the number of PSD95 nanodomains present^43^, and to AMPA receptor (AMPAR) expression and synapse size/strength^41^. Moreover, the number of PSD95 nanodomains at a synapse is proposed to be directly related to the number of AMPAR at a synapse^42,44,47^. Synaptic PSD95 nanodomain number therefore likely predicts synaptic AMPAR expression and in turn, synapse strength, which is determined by the number of AMPAR present. On this basis, the *Der1* mutant mouse neurons are predicted to possess fewer synaptic AMPA receptors and weaker synapses.

AMPAR-induced depolarisation initiates NMDAR-dependent synaptic plasticity and long-term potentiation via molecular pathways involving signalling molecules such as CAMKII, PKA, PDE4 and the transcription factor CREB^48–52^. This leads to increased synaptic AMPAR expression and dendritic spine volume through actin remodelling, and thus greater synapse strength and size^53,54^. Various DISC1 mutant mice exhibit synaptic plasticity defects^34^ and in some cases involvement has been demonstrated of PDE4, which interacts with and is regulated by DISC1^25^ (Millar et al., 2005), or CREB.^34^ Moreover, DISC1 is known to regulate AMPAR subunit expression and dendritic spine size/density downstream of NMDA receptors via actin remodelling^11,55^, and could therefore control plasticity through these mechanisms. DISC1 is also reported to regulate NMDAR activity through PDE4-mediated activation of CREB-dependent GluN2A expression^12^, although we found no evidence of this mechanism in *Der1* mice since hippocampal GluN2A levels are unaltered. DISC1 may therefore regulate synaptic plasticity at multiple levels downstream of NMDAR, and it is notable that the gene expression changes in the human neurons derived from t(1;11) carriers, including altered expression of *PDE4B* and genes required for actin cytoskeleton remodelling are a good fit with these processes. Crucially however, our data indicate that DISC1 has the potential to modulate the triggering of plasticity itself, through effects upon synapse strength and via direct interaction with GluN1, regulation of NMDAR dynamics, and thus control of their synaptic expression and availability to initiate plasticity. Synaptic plasticity underlies cognition which is characteristically impaired in schizophrenia in particular, but also in affective disorders. Our findings thus point towards a pathway that is of direct relevance to a core feature of psychiatric disorders in t(1;11) carriers, and to major mental illness in general.

Four potential genetic modifiers have recently been identified in the t(1;11) family^30^. Of the genes of interest identified at these loci, only *GRM5* and *CNTN5* could be meaningfully examined using the selected panel of t(1;11) family neurons and, although abundantly expressed, neither was found to be dysregulated. Neither were any deleterious sequence variants that would affect protein function discovered in these genes^30^, although potential expression quantitative trait loci (eQTLs) that could affect *GRM5* expression levels were previously identified within the modifier^30^. Such eQTLs may therefore act upon *GRM5* in a cell type-specific manner. The *Der1* mouse mimics disruption of *DISC1* by the t(1;11), isolating this event from any effects of the putative modifiers. We therefore propose that the deleterious effects of DISC1 disruption upon excitatory synapses, identified here using the *Der1* mutant mouse, contribute substantially to risk of major mental illness in t(1;11) carriers, but can be influenced by the genetic modifiers and/or environmental factors, leading to differing outcomes. This is consistent with a previous observation that, irrespective of diagnosis, all tested t(1;11) carriers exhibit an abnormal schizophrenia endophenotype, the P300 event-related potential^6^, and that P300 abnormalities are linked to NMDAR dysfunction^56^.

In conclusion, the *Der1* mutation-induced NMDAR trafficking and synapse abnormalities described here converge upon themes of dysfunctional NMDAR, excitatory synapses and plasticity that are emerging from schizophrenia and depression GWAS and CNV data. The t(1;11) thus provides a mechanistic window into the biochemical basis of these disorders, which links to genetic findings, and highlights new ways to consider therapeutic intervention.

## CONFLICT OF INTEREST

The authors declare no conflict of interest.

## ACKNOWLEDGEMENTS

We thank the t(1;11) family members who have taken part in our research by donating skin biopsies for these experiments. This work was funded by MRC grants G0902166 and MR/J004367/1; Brain & Behavior Research Foundation Independent Investigator Grant 23306; a University of Edinburgh Wellcome Trust Institutional Strategic Support Fund award; European Union Seventh Framework Programme 607616FP7, Deciphering inter- and intracellular signalling in schizophrenia, and a Wellcome Trust Clinical Fellowship to MJ (WT/100135/Z/12/Z).

## SUPPLEMENTARY INFORMATION

**Supplementary information contains detailed materials & methods, Supplementary Figures 1-10 And Supplementary Table 2**

**Supplementary Table 1a-g**

**Supplementary Video** Bidirectional movement of photoconverted GluN1-Dendra2 away from the photoconversion ROI (yellow box in frame 1) in a representative *Disc1*^*w/wt*^ hippocampal neuron. Scale bar, 15μm

## SUPPLEMENTARY INFORMATION

### MATERIALS AND METHODS

#### Generation of induced pluripotent stem cells (IPSC) from dermal fibroblasts

Ethical consent relating to the translocation family is as follows: Prior to 2014, Lothian Research and Development (2011/P/PSY/09), Scotland A Research Ethics Committee (09/MRE00/81); 2014 onwards, Lothian Research and Development (2014/0303), Scotland A Research Ethics Committee (14/SS/0039). Fibroblasts were cultured in DMEM with 10% FBS (all media and supplements from Life Technologies unless stated otherwise) at 37°C with 5 % CO_2_. Fibroblasts were reprogrammed by non-integrating methods, using episomal plasmids^1^. For episomal reprogramming of some lines we used a protocol adapted by Tilo Kunath, (University of Edinburgh) and Roslin Cells (roslincells.com). Other lines were reprogrammed by Roslin Cells. Plasmids incorporating Oct3/4, shRNA to p53, SOX2, KLF4, L-MYC and LIN28 were electroporated into fibroblasts using Nucleofection (Amaxa, Lonza). The episomal plasmids pCXLE-hOCT3/4-shp53-F (for OCT3/4 and p53 knockdown), pCXLE-hSK (for SOX2 and KLF4) and pCXLE-hUL (for L-MYC and LIN28) were a gift from Shinya Yamanaka. (These correspond to Addgene plasmids 27077, 27078, 27080). 5 × 10^5^ fibroblasts were transfected with 1.7ug of pCXLE-hOCT3/4-shp53-F, 1.6ug of pCXLE-hSK and 1.7ug of pCXLE-hUL using the NHDF Nucleofector kit and Amaxa Nucleofector protocol U-023 (1,650 V, 10 ms, 3 time pulses), according to the manufacturer’s instructions. Cells were then seeded into one well of a gelatin-coated 6 well tissue culture grade plate in Opti-MEM (ThermoFisher Scientific) supplemented with 10% FCS and 1% Antibiotic Antimycotic Solution (Life Technologies 15240062). All stages of cells were maintained in media supplemented with Antibiotic Antimycotic Solution thereafter. The medium was replaced every 2-3 days before cells were replated into a 10cm Vitronectin/Geltrex (ThermoFisher Scientific) or Matrigel (Life Technologies)-coated tissue culture grade dish after 6-7 days. The following day the medium was changed to Essential 6 medium (ThermoFisher Scientific) with added 100 ng/ml bFGF (Peprotech). The medium was changed every 2 days until colonies were ready to be picked, at approximately day 25-30. Individual colonies were picked and expanded into 12, then 6, well Vitronectin/Geltrex or Matrigel-coated tissue culture grade plates in Essential 8 medium (ThermoFisher Scientific) with daily medium changes. Cells were passaged using 0.5mM EDTA in PBS. IPSCs were generated and cultured at 37°C, 5% CO_2_ and 21% O_2_. Quality control of IPSC lines was performed after clonal passage 10.

Pluripotency of IPSC lines was assessed using markers SSEA-1 PE, SSEA-4-AlexaFluor647 and Oct3/4 PerCP-Cy5.5 and isotype controls using the BD Stemflow Human and Mouse Pluripotent Stem Cell Analysis Kit (BD Biosciences, 560477) according to the manufacturer’s instructions, or with SSEA-1-APC (301907), SSEA-4-FITC (330409), TRA-1-60-PE (330609), TRA-1-81-PE (330707) and isotype controls (all BioLegend) as follows: IPSCs were dissociated using StemPro Accutase (Life Technologies) and washed with Essential 8 medium. 1×10^5^ cells were incubated with antibodies in 2% Fetal Bovine Serum in PBS for 1 hour on ice. Cells were washed once with 2% BSA/PBS, centrifuged at 200 × g for 5 min and resuspended in 200 ul 2% BSA/PBS. FACS analysis was performed on single cell suspensions using a FACS Aria cell sorter (BD Biosciences). Data were analysed using FlowJo v10 software.

EBNA-1 primer sequences and amplification protocol were taken from the Epi5 Episomal iPSC Reprogramming Kit (Life Technologies, A15960). Genomic DNA was extracted from IPSCs using the DNeasy kit (Qiagen). Cells that had not been in contact with episomes, were used as negative controls. Positive controls were low passage IPSC lines where episomes were still present. A non-template control (NTC) was also used. IPSC lines were only taken forward once episomal clearance had been confirmed by this method (data not shown).

The human Cytoscan 750 K Array (Affymetrix) was used to identify genomic abnormalities in the IPSC lines. The array consists of 550,000 unique non-polymorphic probes and 200,000 SNPs for accurate genotyping. Genomic DNA was extracted from IPSC clones using the DNeasy kit (Qiagen) according to manufacturer’s instructions. Samples were sent to the NHS Cytogenetics Laboratory (Western General Hospital, Edinburgh) for processing of the arrays. Chromosome analysis was performed using Chromosome Analysis Suite version 2.0 (Affymetrix). Copy number, breakpoints and Loss Of Heterozygosity (LOH) regions were determined using the models and algorithms incorporated within the software package. To exclude possible false positives due to inherent microarray noise the CNV threshold of gains and losses for inclusion in analyses was 10 kilobase pairs (kbp) and 10 consecutive markers. IPSC lines with deletions or duplications greater than 5 MB, the limit typically applied by G-banding, were excluded from further studies.

#### NPC culture and neuronal differentiation

IPSCs were converted into neuroectoderm by dual-SMAD signalling inhibition^2^. Long-term anterior neural precursor cells were generated and maintained under physiological normoxia (3% O2) and in the absence of EGF^3^. NPCs were cultured at 37°C, with 5% CO_2_ and 3% O_2_ on Matrigel (Life Technologies)-coated 6 well tissue culture grade plates in Advanced DMEM/F-12 (Life Technologies) with 1% Glutamax-1 (Life Technologies), 1% N2 supplement (Life Technologies), 0.1% B27 supplement (Life Technologies), 10 ng/ml bFGF (PeproTech) and 1% antibiotic/antimycotic solution (Life Technologies). NPCs were maintained up to passage 30 with feeding every 2-3 days and weekly passages using StemPro Accutase (Life Technologies). All NPC lines were tested every week for mycoplasma infection. For differentiation into cortical forebrain-like neurons^3^, NPCs were plated into Matrigel (Life Technologies)- and Laminin (Sigma-Aldrich)-coated 12 well tissue culture grade plates in Advanced DMEM/F-12 with 0.5% Glutamax-1, 0.5% N2 supplement, 0.2% B27 supplement, 2 μg/ml Heparin and 1% antibiotic/antimycotic solution (Life Technologies). Neurons were maintained for 5 weeks with feeding as necessary. During weeks 2 and 3 the neuronal differentiation medium was supplemented with Forskolin (Tocris Bioscience). During weeks 4 and 5 the Forskolin was removed, and the medium was supplemented with BDNF (Life Technologies) plus GDNF (Life Technologies) to 5ng/ml each.

#### Mouse generation and colony maintenance

VelociMouse^®^ technology (Regeneron)^4^ was used to target embryonic stem cells and microinject them into mouse embryos. In brief, F1H4 (129S6SvEv/C57BL6F1) embryonic stem cells were electroporated with the linearized vector construct and positive clones were microinjected into 8-cell stage mouse C57BL6 embryos. Microinjected embryos were transferred to uteri of pseudopregnant recipient females, weaned pups were scored, and high percentage chimera males were selected for mating with flp-positive C57BL6 females to remove the selection cassette, to prove germ-line transmission, and to generate F1 animals for further breeding.

Because there is already a mutation (25bp deletion) at the *Disc1* allele in exon 6 in the 129/Sv strain which causes a truncation of *Disc1*^5^, F1 progeny were generated and a PCR assay which distinguishes the C57BL/6 allele versus the 129/Sv allele was employed to determine which F0 mice were correctly targeted to the C57BL/6 locus (Supplementary Figure 10). Mice which carried the translocation on the C57BL/6 allele were then crossed to CMV-Cre mice to remove the Neo cassette via Cre-mediated recombination at the flanking loxP sites. Genotyping results were confirmed by Loss-of-Native-Allele assay.

The exclusion of the differentially spliced *DISC1FP1* exon 3a^6^ that is present in a minority of transcripts (www.genome.ucsc.edu) precludes production of transcripts encoding CP1. The exclusion of the differentially spliced *DISC1FP1* exon 7b does not affect the potential production of CP60/69 proteins since the stop codon in chimeric transcripts encoding these proteins occurs in exon 6^6^. Since the *Disc1* allele was modified on a mixed background of 129 and C57BL/6J, a congenic breeding strategy was adopted to purify the strain background. Following repeated crossing to C57BL/6J mice, genotyping of polymorphic markers carried out by the Jackson Laboratory found the mice to be >99.5% C57BL/6J. These mice were then mated to C57BL/6J for one final round and the progeny used for subsequent experiments. Mice were housed in the Biomedical Research Facility at the University of Edinburgh. All mice were maintained in accordance with Home Office regulations, and all protocols were approved by the local ethics committee of the University of Edinburgh.

#### Expression constructs

Constructs pCB6-HA-GluN2B^7^, expressing full-length rat GluN2B tagged at the N terminus with the HA epitope, pCB6-GluN1^7^ and a plasmid expressing HA-GluN1 were gifts from William Green (University of Chicago). A GluN1-YFP expression construct was a gift from Seth Grant (University of Edinburgh).

Full length GluN1 and the C-terminal tail of GluN1 (C0-C1-C2) were subcloned into pCMV4A (Agilent) to generate FLAG-tagged constructs using the following Not1-Sal1 primer combinations: gatcgcggccgcaccatgagcaccatgcacctgctgacattcgcc, gatcgtcgacgctctccctatgacgggaacac and gatcgcggccgccaccatggagatcgcctacaagcgacacaag, gatcgtcgacgctctccctatgacgggaacacagctg. The GluN1 C0-C1-C2 sequence was subcloned into the BamH1-Not1 site of pEBG-2T to produce a GST fusion.

A DISC1 open reading frame consisting of amino acids 358-499 was inserted into the BamH1-Not1 sites of pEBG to generate pEBG GST-DISC1 358-499. The full-length DISC1 open reading frame was inserted into the same plasmid to generate pEBG GST-DISC1.

Plasmid pGluN1-Dendra2, expressing full-length human GluN1 (transcript variant GluN1-1a) tagged at the COOH terminus with Dendra2, was generated as follows. Dendra2 coding sequence was amplified from pDendra2-N (Clontech) using primer pair gatcgcggccgctcgagatgaacaccccgggaattaacc and gatcggccggccttaccacacctggctggg and sub-cloned between the NotI and FseI sites of construct pCMV6-AC-NR1 (GluN1)-GFP (Origene RG216458).

Generation of pcDNA4/TO constructs expressing FLAG-TRAK1, FLAG-DISC1, FLAG-DISC1-37W and a corresponding empty vector has been described previously^8, 9^. All expression constructs were verified by sequencing.

#### RNA isolation and cDNA synthesis

Total RNA was extracted using the RNeasy mini Kit (Qiagen), according to the manufacturer’s instructions. Genomic DNA was removed prior to cDNA synthesis using the TURBO DNA-free Kit (Thermo Fisher Scientific) and cDNA synthesis was performed using Dynamo cDNA synthesis kit (Thermo Scientific).

#### RT-PCR

RT-PCR was performed using the Titanium Taq PCR Kit (Clontech). Each reaction contained cDNA template, 2.5μl PCR buffer, 0.5μl dNTPs, 1μl of both primers (10μM stock) and 0.5μl enzyme in a total volume of 25μl. Non-template controls were created by replacing the cDNA template with water.

#### Quantitative RT-PCR

Quantitative Real-Time PCR was performed in 384-well plates using Power SYBR green PCR Master Mix (Applied Biosystems) on the 7900 HT sequence Detention System (Applied Biosystems). Non-template and minus reverse transcriptase controls were included in all experiments with three technical replicates for all samples. To control for inter-plate variation a calibrator sample was included on every plate for normalisation purposes. For quantification of human DISC1, housekeeping gene stability was assessed across samples taken from NPCs through to five week neurons for several genes using geNorm (genorm.cmgg.be/). *GAPDH* and *ACTB* were subsequently selected as the most stable housekeeping genes for use in quantitative RT-PCR in these samples. For quantification of mouse *Disc1*, *Cyclophilin* and *Hmbs* were found to be stable in the mouse brain samples analysed. *ACTB* was used as a reference gene for neuron RNASeq follow-up. Primer pairs were optimised for amplification efficiency and their specificity confirmed by PCR product sequencing. Melting curve analysis was carried out for each primer pair to optimise amplification conditions and confirm amplification specificity. PCR efficiency was assessed by running standard curves using serial dilutions of NPC or mouse brain samples for human and mouse primers, respectively. Gene expression levels were calculated using the relative standard curve method, with normalisation to the geometric mean of the reference genes

A typical reaction contained 5μl of Master Mix, 0.6μl of both primers (10μM stock) or 0.2μl in the case of *Hmbs* primers and 4μl of 1/10 diluted cDNA to a total volume of 10μl. Amplification was achieved in 40 cycles of the following conditions: 50°C for 2min, 95°C for 10min, 95°C for 15 sec 65°C for 45 sec, followed by the dissociation curve: 95 °C for 15 sec, 60°C for 15 sec, 95 °C for 15 sec.

#### PCR primer sequences

Human RT-PCR primers were as follows:

derived 1 CP1 chimeric transcript F (*DISC1* exon 6/7): aaggagcctccaggaaagaa
derived 1 CP1 chimeric transcript R (*DISC1FP1* exon 3a): caagaaatgccaaagtgagtt
derived 1 chimeric transcripts pan F (*DISC1* exon 6/7): aaggagcctccaggaaagaa
derived 1 chimeric transcripts pan R (*DISC1FP1* exon 4): aaggagcctccaggaaagaa
derived 11 chimeric transcripts F (*DISC1FP1* exon 2): gggacctggaattgaagaga
derived 11 chimeric transcripts R (*DISC1* exon 9): gtctcctggtgctccacttc
*DISC2* F: ccctgaaggtgttgaacaagc
*DISC2* R: ctggaccctctgttgctgta
*DISC1FP1* F: agagcaagaagagtggatgtgga
*DISC1FP1* R: ccttgaggagtacgtcttaagctct

Human RT-QPCR primers were as follows:

*DISC1* F: ccagccttgcttgaagccaaaa
*DISC1* R: tgaggagtccctccagcccttc
*GAPDH* F: gagtccactggcgtcttcac
*GAPDH* R: atgacgaacatgggggcatc1
*ACTB* F: gttacaggaagtcccttgccatcc
*ACTB* R: cacctcccctgtgtggacttggg
*ERBB4* F: agagcccaccaattactcca
*ERBB4* R: gtgtaacggtcccactagtca
*PDE4B* F: cacggcgatgacttgattgt
*PDE4B* R: tgtgggttgactctggagac
*NRG1* F: aacaaagcatcactggctga
*NRG1* R: aagacacatatgctccttcagttg
*DRD2* F: aagggcacgtagaaggagac
*DRD2* R: ggtcaccgtcatgatctcca
*GRM5* F: ggagcttgattgtgatgcca
*GRM5* R: tgcttctgtgagggcatga
*CNTN5* F: tcaggcggtgctggaaata
*CNTN5* R: ggctcccactgtctaactga

Mouse RT-PCR primers were as follows:

derived 1 chimeric transcripts pan F (*Disc1* exon 8): gtgctcaggtgagaagctgtg
derived 1 chimeric transcripts pan R (*DISC1FP1* exon 4): gacccacagatggaatcgaa

Mouse RT-QPCR primers were as follows:

*Disc1* F: cctgccttgctggaagcca
*Disc1* R: cccttcccgctctgacgaca
Derived 1 chimeric transcripts pan F (*Disc1* exon 8): cctgccttgctggaagcca
Derived 1 chimeric transcripts pan R (*DISC1FP1* exon 4): aagacccacagatggaatcgaactg
*Cyclophilin* F: ggagatggcacaggaggaaag
*Cyclophilin* R: gcccgtagtgcttcagcttgaa
*Hmbs* F: ccctgaaggatgtgcctaccata
*Hmbs* R: aaggtttccagggtctttccaa

Mouse genotyping was carried out using a multiplex reaction with the following primers:

*Disc1* F (intergenic sequence beyond final exon): cctgcatccacagacgtgc
*Disc1* R (intergenic sequence beyond final exon): cagtagtaagaaaagagacaaccccc
Targeting vector F: ataacggtcctaaggtagcgagc

#### PCR product sequencing

Where single PCR products were obtained, completed PCR reactions were treated with ExoSAP-IT (GE healthcare) prior to direct sequencing. Alternatively, where multiple products were generated (Supplementary Figure 3c), PCR products were excised from agarose gels and purified using the QIAquick Gel Extraction Kit (Qiagen). BigDye Terminator sequencing of PCR products was carried out using 1 μl BDv3.1, 1.5 μl 5X sequencing buffer, 1 μl DNA template and 6 μl dH_2_O. Reactions were cycled as follows; 96°C-1 min (1 cycle); 96°C-10 sec, 50°C-5 sec, 60°C-4 min (25 cycles); 4°C hold. Sequencing chromatograms were analysed using Chromas (version 1.45).

#### RNA sequencing

High density neuronal cultures (approximately 2 million cells per well in 12 well plates) were differentiated for 5 weeks. Immunofluorescence staining of parallel cultures was used to confirm correct NPC morphology and Nestin expression at the time of plating, and successful neuronal differentiation by assessing morphology and acquisition of βIII-tubulin expression at the time of harvesting, for every culture used. Three independent neuronal differentiations were performed per NPC line. Neurons were harvested in RNAlater (ThermoFisher Scientific), stored at −80°C, then processed in batches to extract the RNA. Each batch consisted of one triplicate per line to minimise batch effects.

All subsequent steps were performed by the Wellcome Trust Edinburgh Clinical Research Facility (www.wtcrf.ed.ac.uk). Total RNA samples were assessed on the Agilent Bioanalyser (Agilent Technologies, G2939AA) with the RNA 6000 Nano Kit (5067-1511) for quality and integrity of total RNA, and then quantified using the Qubit 2.0 Fluorometer (Thermo Fisher Scientific Inc, Q32866) and the Qubit RNA BR assay kit (Q10210). Samples were also assessed for DNA contamination using the Qubit DNA HS assay Kit (catalogue Q32851).

Libraries were prepared from each total-RNA sample using the TruSeq Stranded Total RNA with Ribo-Zero Gold kit (RS-122-2301) according to the provided protocol. 500ng of total-RNA was processed to deplete rRNA before being purified, fragmented and primed with random hexamers. Primed RNA fragments were reverse transcribed into first strand cDNA using reverse transcriptase and random primers. RNA templates were removed and a replacement strand synthesised incorporating dUTP in place of dTTP to generate double-stranded cDNA. AMPure XP beads (Beckman Coulter, A63881) were then used to separate the double-stranded cDNA from the second strand reaction mix, providing blunt-ended cDNA. A single ‘A’ nucleotide was added to the 3’ ends of the blunt fragments to prevent them from ligating to another during the subsequent adapter ligation reaction, and a corresponding single ‘T’ nucleotide on the 3’ end of the adapter provided a complementary overhang for ligating the adapter to the fragment. Multiple indexing adapters were then ligated to the ends of the double-stranded cDNA to prepare them for hybridisation onto a flow cell, before 15 cycles of PCR were used to selectively enrich those DNA fragments that had adapter molecules on both ends and amplify the amount of DNA in the library suitable for sequencing.

Libraries were quantified by PCR using the Kapa Universal Illumina Library Quantification kit complete kit (KK4824) and assessed for quality using the Agilent Bioanalyser with the DNA HS Kit (5067-4626). Libraries were combined in three equimolar pools and sequencing was performed using the NextSeq 500/550 High-Output v2 (150 cycle) Kit (FC-404-2002) on the NextSeq 550 platform (Illumina Inc. SY-415-1002). Sequences were aligned to the human reference genome Hg19 using the RNA-Seq Alignment v1.0 application (Illumina Inc.).

#### RNA sequencing data analysis

Differential gene expression was analysed using DESeq2 from the R statistical package^10^. Differential exon expression was analysed using DEXSeq^11^. The three replicate samples used per cell line were obtained from independent neuronal differentiations which can follow slightly different trajectories. These replicates were therefore treated as biological rather than technical replicates. Due to sex imbalance in the samples all differentially expressed genes from the X and Y chromosomes were removed from the DESeq2 and DEXSeq analysis. For all downstream analyses, the full list of expressed genes was used as the background gene set, while an expression BaseMean cut-off of ten was applied to all differentially expressed data (corrected p<0.05) to minimise quantitation errors from genes expressed at very low levels. DESeq2 and DEXSeq data were combined and GO analysis was carried out using Gorilla (http://cbl-gorilla.cs.technion.ac.il/). Enrichment of GO terms categorised under Process, Function and Component was investigated. Ingenuity Pathway Analysis was carried out to complement the gene ontology analysis. Heat maps of gene expression were generated using R version 3.4.2 and RStudio version 1.0.143. Raw count data for all samples were together subjected to a regularised logarithm transformation^10^ using the DESeq2 package version 1.16.1. For each heat map, the transformed counts for each gene were normalised to Z-scores across all samples and subsequently visualised using the pheatmap package version 1.0.8 (cran.r-project.org/package=pheatmap).

#### Peptide arrays

Peptide libraries were produced by automatic spot synthesis as described previously^12^, ^13^. Interaction of peptide spots with purified proteins was determined by overlaying the cellulose membranes with 10μg/ml recombinant protein. Membranes were incubated with protein that had been partially purified from 500μl of reticulocyte lysate and bound protein was detected by immunoblotting using specific primary antisera and a complementary HRP-coupled secondary antibody. For alanine scanning the peptide was systematically mutated to alanine at every position, except existing alanine residues which were mutated to aspartate.

#### *In vitro* transcription and translation

FLAG-DISC1 and FLAG-GluN1 C0-C1-C2 proteins were synthesized using the T3/T7 transcription/translation-coupled reticulocyte lysate system (Promega, Hampshire, UK). The translation product was partially purified by ammonium sulphate precipitation before use in peptide array analyses.

#### Cell and tissue lysis

Cultured cells were lysed in ice-cold PBS containing 1% Triton X-100/10mM sodium fluoride/1mM DTT/2mM PMSF/5mM pyrophosphate/10% glycerol containing protease inhibitor cocktail (Roche) and phosphatase inhibitor cocktail II (Calbiochem). Lysates were solubilised by incubation for 30-60 minutes at 4°C on a rotary wheel and centrifuged at 13,000 rpm for 30min. Brain lysates were prepared from dissected tissue and homogenised in ice-cold PBS containing 1% Triton X-100/1% NP40/0.5% sodium deoxycholate/10% glycerol/1 mM DTT/ 10 mM β-glycerophosphate/10 mM NaF/phosphatase inhibitor cocktail II and 1V (Calbiochem) and 2X protease inhibitor cocktail (Roche), solubilised by incubation on a rotary wheel for 30 mins at 4°C and centrifuged at 45,000 rpm for 30 mins at 4°C to obtain the soluble fraction.

#### Immunoblotting

Protein samples were separated by SDS-PAGE using NuPAGE polyacrylamide gels (Invitrogen), transferred onto PVDF membranes (GE Healthcare) using Trans-Blot SD SemiDry Transfer Cell (Bio-Rad) and blocked in 1% skimmed milk in T-TBS [50 mM Tris-HCl (pH 7.5), 150 mM NaCl and 0.1% Tween-20] at room temperature (RT). Incubation with primary antibodies was performed overnight at 4 °C. Blots were incubated with horseradish peroxidase-conjugated secondary antibodies [Rabbit anti-mouse IgG HRP, Swine anti-rabbit IgG HRP or Rabbit anti-goat IgG HRP (DAKO)] for 45 min at RT and then protein bands were visualized by ECL or ECL-2 Western blotting substrate (Thermo Scientific). Some blots were stripped with Restore PLUS Western Blot Stripping Buffer (Thermo Scientific) and reprobed with appropriate antibodies, including secondary antibodies conjugated to alkaline phosphatase, with protein bands subsequently visualized using Western Blue (Promega). Relative intensities of protein bands were quantified with ImageJ densitometry analysis.

#### Subcellular Fractionation of Mouse Brain

Crude synaptosomes and light membrane fractions prepared as described previously^14^ have already been reported^15^. Briefly, C57BL/6 adult mouse brains were homogenized in a sucrose buffer solution [1mM HEPES (pH7.5), 0.32M sucrose, 1mM NaHCO_3_, 1mM MgCl_2_ Complete Mini Protease Inhibitor Tablet (Roche), Phosphatase Inhibitor Cocktail Set II (Calbiochem)] using a dounce tissue grinder. The S1 supernatant was obtained by centrifugation of the homogenates at 1,000*g* for 10 min and further fractionated into the S2 supernatant and the P2 crude synaptosomal pellets by centrifugation at 13,800 *g* for 10 min. To obtain the light membrane P3 pellets, the S2 supernatant was centrifuged at 100,000 *g* for 1 hr (Beckman TLA 100.3). The identity of the synaptosome fraction was confirmed previously by probing for PSD95^15^.

#### Immunoprecipitation

Cultured cells were lysed in IP buffer [1% Triton X-100, with/without 0.1% SDS, 50 mM Tris-HCl (pH 7.5), 150 mM NaCl, Complete Mini Protease Inhibitor Tablet (Roche), Phosphatase Inhibitor Cocktail Set II (Calbiochem)]. Insoluble materials were removed by centrifugation at 100,000g. Crude synaptosomes/light membrane pellets were lysed in IP buffer and insoluble materials were removed by ultracentrifugation at 100,000g (Beckman TLA 100.3) for 30 min at 4°C. Cell lysates were pre-cleared with protein G-Sepharose beads (Sigma) alone for 1hr at 4°C, and then incubated with appropriate antibodies overnight at 4°C with rotation. Protein G-Sepharose beads were added to the lysates and further incubated for 2 hrs. Immune complexes on the beads were washed with IP buffer three times and with 50mM Tris-HCl (pH 7.5) once. Bound proteins were eluted with SDS loading buffer and analyzed by immunoblotting.

#### Antibodies

A new antibody specific for human DISC1 was generated as described previously^15^. Briefly, the C-terminus of human DISC1 isoform Lv (aa 669-832) was bacterially expressed as a GST-fusion protein and purified using Glutathione Sepharose 4B (GE Healthcare). New Zealand White rabbits were then immunized with the GST-human DISC1 669-832 protein and immune sera were collected (Eurogentec). Anti-DISC1 polyclonal antibodies were purified using Activated CH Sepharose 4B (GE Healthcare) to which the same antigen was covalently coupled. Antibodies that reacted with GST were removed from the immune sera using GST-coupled Activated CH Sepharose 4B prior to the affinity purification of anti-DISC1 antibodies. Generation and characterisation of an in-house antibody to mouse Disc1 has previously been described^15^, ^16^.

The following antibodies were used in the surface NMDAR expression study: GluN1 Mouse monoclonal IgG2a (BD Biosciences, 556308), GluN2A Rabbit polyclonal (Alomone Labs, AGC-002), GluN2B Rabbit polyclonal (Alomone Labs, AGC-003), βIII-Tubulin Rabbit polyclonal (Abcam, Ab18207), βIII-Tubulin/Tuj1 Mouse monoclonal IgG2a (Cambridge Bioscience, 801201) and PSD95 Mouse monoclonal IgG1k (ABR, MA1-046). Fluorescently labelled secondary antibodies were all AlexaFluor (Life Technologies): Goat anti mouse IgG1 488, Goat anti mouse IgG2a 647 and 594, Chicken anti rabbit 647, Goat anti rabbit 594.

The following antibodies were also used in this study: mouse monoclonal anti-FLAG M2 (Sigma F3165), anti-rabbit NR1 (GluN1, Sigma 8913), goat anti-HA (Abcam ab9134), rabbit anti-Calreticulin (Sigma 06-661), mouse monoclonal anti-Vinculin (Abcam ab18058) and mouse monoclonal anti-Nestin (Millipore 10C2).

#### Cell culture and transfection

All cultures were maintained at 37 °C and 5% CO_2_ in a humidified incubator. COS7 cells were grown in DMEM/10% foetal calf serum and passaged using a 1:1 trypsin:versene solution (Invitrogen). COS7 cells were transfected at 60-70% confluency using Lipofectamine 2000 (Life Technologies) according to the manufacturer’s instructions.

Hippocampal neurons were isolated from embryonic (E18) C57BL/6 mouse brains as described previously^9^, and grown in the presence of an astrocyte feeder layer. Astrocytes were purified from E18 C57BL/6 mouse brains as described previously^17^ and grown to confluence on rat tail collagen coated 30mm PTFE cell culture inserts (Millipore) in DMEM with GlutaMAX (Life Technologies) supplemented with 10% FCS. The astrocyte-containing cell culture inserts were then transferred to Poly-D-Lysine coated 6 well high precision (1.5H) glass bottom plates (In Vitro Scientific), and the medium was replaced with Neurobasal medium without phenol red supplemented with B27 and GlutaMAX (Life Technologies). Two days later, hippocampal neurons were seeded at low density (1×10^5^/well) underneath each astrocyte-containing cell culture insert and in the presence of the astrocyte-preconditioned medium. When cultures were grown for longer than one week, the entire volume of medium (2 ml) in the astrocyte-containing insert was refreshed one week after plating, and every subsequent week afterwards. The genotype of each litter used to isolate hippocampal neurons was confirmed by PCR using the tail tips of the individual embryos.

For live imaging of GluN1-Dendra2 trafficking in dendrites, neurons were transfected at DIV7 with Lipofectamine 2000 (Life Technologies). A total of 1.75μg of endotoxin-free plasmid DNA (750 ng of pGluN1-Dendra2 plus 1μg of pCB6-HA-GluN2B) and 2μl of Lipofectamine 2000 were used per well. Plasmid DNA and Lipofectamine2000 were diluted in 50μl each of Neurobasal medium without phenol red, mixed thoroughly and incubated at 37 °C for 1 hour. 400μl of medium were removed from the astrocyte-containing insert located in the well to be transfected, and used to dilute the transfection mix. The astrocyte-containing insert was then temporarily removed from the well, the entire volume of medium in the well was set aside and replaced with the diluted transfection mix, and then the astrocyte insert was returned to the well. After incubating for 4 hours at 37 °C and 5% CO_2_, the transfection mix was removed, and the pre-existing Neurobasal medium was returned to the well. The astrocyte insert was left in place, and replenished with fresh Neurobasal medium without phenol red supplemented with B27 and GlutaMAX. The transfected neurons were imaged on the day following transfection.

#### Dendra2 NMDAR trafficking assay

Monomeric fluorescent protein Dendra2 exhibits green fluorescence (excitation/emission peaks: 490 nm/507nm) in its native state, and can be stably photoconverted to red fluorescence (excitation/emission peaks: 553nm/ 573nm) by exposure to intense 405nm light. Because GluN2 subunit availability is a limiting factor for receptor assembly^18^, GluN1-Dendra2-expressing neurons were co-transfected with an HA-GluN2B expression construct to maximise NMDAR assembly, and therefore trafficking outwith the ER.

C57/BL/6 hippocampal neurons expressing GluN1-Dendra2 plus HA-GluN2B were imaged at DIV8 (one day after transfection) using a Nikon C1 confocal microscope equipped with a Plan Apo 60x oil immersion objective (NA 1.4), and an environmental chamber. Neurons were imaged in their original culture medium, and in the presence of the astrocyte-containing insert. Temperature and CO_2_ saturation were kept constant at 37°C and 5%, respectively, throughout the imaging period. To locate transfected neurons, the sample was observed with a FITC filter, using the minimum light intensity that still allowed green fluorescent signal to be obtained. This was necessary to prevent unwanted bulk Dendra2 photoconversion. After switching to confocal mode, the zoom factor and focus were rapidly adjusted while exciting the sample with a 488nm laser at low power (1%). To verify that no unwanted photoconversion had occurred, an image of the same field of view was then obtained in the red channel by exciting the sample with a 561nm laser, confirming that no signal above background was detectable.

Transfected primary dendrites expressing GluN1-Dendra2 were selected and a small rectangular region of interest (ROI) was positioned in a central location along its longitudinal axis (Supplementary Figure 5c). This ROI was then used to target the 405nm laser to induce GluN1-Dendra2 photoconversion. The exact position and size of this ROI (termed the photoconversion ROI) was adjusted for each dendrite based on its size, thickness and brightness, with the aim of obtaining a sufficient amount of photoconverted protein to allow successful tracing. The average length along the dendrite of the photoconversion ROI was ~14μm, and did not vary significantly between experimental groups, ensuring comparability between measurements.

GluN1-Dendra2 photoconversion was induced by rapidly scanning a suitably positioned dendritic ROI with a 405nm laser (laser power: 10%, loops per second: 3, total stimulation time: 1 second, Supplementary Figure 5d). Immediately after photoconversion, acquisition of a time series was started manually (resulting in an effective delay of 10 seconds between photoconversion and acquisition of the first image frame). The time series consisted of 13 frames acquired at 15 second intervals, for a total imaging time of 180 seconds. This frame rate and total acquisition time were determined empirically, and were found to be the best compromise between fluorescence preservation over time (a higher frame rate would have induced more rapid photobleaching) and sufficiently accurate documentation of protein movement. The first frame contained both red (photoconverted GluN1-Dendra2) and green (native GluN1-Dendra2) channels, and all subsequent frames contained the red channel only, allowing movement of photoconverted GluN1-Dendra2 to be traced along the dendrite. Acquisition settings were as follows, and were kept consistent throughout the experiment. Frame size: 512 × 512 pixels, scan speed: 1 frame per second, pinhole size: 150μm (fully open), binning: 1×1, focus stabilisation: ON. To ensure comparability between images, laser power and detector settings were kept consistent throughout the experiment.

To quantify the movement of photoconverted GluN1 signal along the dendrite, a bespoke analysis algorithm was generated using iVision software (BioVision Technologies, script available on request). This algorithm uses the green (non-photoconverted) image of GluN1-Dendra2 to define the structure of the dendrite to be analysed (Supplementary Figure 5e-j). The resulting ‘skeleton’ corresponding to the longitudinal axis of the primary dendrite and all its branches is superimposed onto the red image, providing a guide or “track” along which the algorithm will then quantify the intensity of the red pixels. Such quantification is achieved by means of a circular analysis ROI whose centre moves along the track by incremental steps corresponding to one radius at a time. Since the data are deleted from the image after each measurement is taken, and before the analysis circle proceeds to the next location, the equivalent of half the area of this circle is analysed at each step, and no pixels are measured twice. The centre of the analysis circle is initially positioned in a location corresponding to the pixel coordinates of the geometric centre of the photoconconversion ROI. After measuring the intensity of the red pixels located within the central dendrite segment (or central bin), the first analysis circle moves along the track, starting from the proximal direction (towards the neuronal soma) and continuing until all proximal dendrite segments (bins) have been analysed through the time-lapse series. Next, the analysis circle repositions itself on the centre of the photoconversion ROI and then moves from here in the distal direction (away from the neuronal soma), proceeding along the distal tracks until all remaining dendritic segments have been analysed through the time-lapse series. Finally, two separate results tables are assembled for the proximal and distal sections of the dendrite. The dendritic segment analysed at each step by the analysis circle is defined by its distance from the centre of the photoconversion ROI, its area, and the sum of the intensities of all red pixels measured within it throughout the time-lapse series. To correct for the effect of variations in dendrite size, total fluorescence intensity was divided by the area of the same segment. For each measured dendritic segment the width of each analysed dendritic segment corresponds to the radius of the analysis circle, which was arbitrarily set to 5 μm. The 5μm bin therefore represents 5-10μm from the photoconversion ROI, and so on. Since red fluorescence intensity did not change significantly over time in bins located beyond 40μm from the centre of the photoconversion ROI (data not shown), only bins located up to 40μm from the centre of the photoconversion ROI were analysed further.

A total of 37, 35 and 40 neurons were imaged per DISC1 expression construct (empty vector, FLAG-DISC1 or FLAG-DISC1-37W, respectively), each from three independent cultures. A total of 69 (*Disc1*^*wt/wt*^) and 68 (*Disc1*^*Der1/Der1*^) neurons were imaged per genotype from three independent cultures. Imaging and analysis was carried out blind to genotype.

Although several neurons were imaged per genotype, the number of neurons that produced usable data decreased in the furthest bins due to a combination of factors. First, not all neurons analysed had dendrites extending at least 40μm on both sides of the photoactivation centre. Second, some statistical outliers (described in SI under were excluded. Third, for each neuron, bins in which there was no clear relationship between fluorescence intensity and time were excluded from the analysis. This was determined by individually plotting their values. Three main classes of fluorescence intensity plot were obtained, corresponding to neurons in which 1) the fluorescence intensity peak was reached within the 10s preceding imaging onset (Supplementary Figure 5k), 2) the peak was reached within the bin (Supplementary Figure 5l), and 3) fluorescence intensity increased throughout the imaging period, most likely due to the peak moving too slowly to be captured within three minutes (Supplementary Figure 5m).

#### Code availability

The bespoke GluN1-Dendra2 motility analysis algorithm generated using iVision software (BioVision Technologies) is available on request.

#### NMDAR velocity estimates

Average velocities were estimated for the slow and fast NMDAR populations, taking into account the approximately 10s delay between photoconversion and imaging onset.

Fluorescence peak average velocities were determined from individual neuron fluorescence intensity plots as follows: The time to reach peak fluorescence was identified for each neuron per condition from the 10μm and 15μm bins. In neurons where fluorescence peaked in the 10s prior to imaging onset (Supplementary Figure 5k) peak fluorescence was set at 10s. All of these values were averaged per bin for each condition. Velocity estimates per bin per condition were then determined by dividing distance between the centre of the photoconversion ROI and the centre of the bin (for example, for the 10μm bin distance was 12.5μm), by time. The 10μm and 15μm bins were used for this analysis because the time to peak fluorescence could be determined for the majority of neurons in these bins, whereas in the 20μm bin, fluorescence did not peak in some neurons, thus any velocity estimates from this bin would be overestimated.

Maximum velocity estimates for the fast NMDAR population were determined from individual neuron fluorescence intensity plots as follows: Neurons from the 25μm, 30μm, 35μm and 40μm bins were examined. In these bins the majority of neurons exhibited a sigmoidal pattern of fluorescence intensity (Supplementary Figure 5n) interpreted as showing that fluorescence appeared in the bin after imaging onset. For these neurons a sigmoidal curve was fitted to the fluorescence intensity plot and the time at which fluorescence intensity first appeared on the linear part of the curve was taken as the time at which fluorescence reached the bin. In these bins a minority of neurons exhibited a pattern of steadily increasing fluorescence (Supplementary Figure 5m). In these neurons it was assumed that fluorescence had already reached the bin by imaging onset, thus fluorescence appearance in these neurons was set at 10s. All of these values were averaged per bin for each condition. Velocity estimates per bin per condition were then determined by dividing distance between the centre of the photoconversion ROI and the centre of the bin in question, by time.

#### Immunocytochemistry

COS7 cells were fixed in 4% Paraformaldehyde (PFA) for 10 minutes and permeabilised with 0.2% Triton X-100 in PBS for 10 minutes. Blocking was carried out for 30 minutes in 3% BSA in PBS, followed by incubation with primary, then secondary, antibodies diluted in 3% BSA/PBS for 1 hour. All steps were carried out at room temperature.

To stain surface-expressed NMDAR on cultured hippocampal neurons, antibodies that selectively bind to the extracellular loop of the receptor subunits (either GluN1 Mouse monoclonal IgG2a, GluN2A Rabbit polyclonal or GluN2B Rabbit polyclonal) were directly added to the culture medium at a dilution of 1:100, and the living neurons were incubated at 37 °C for 10 minutes in a humidified incubator with 5% CO_2_. Neurons were then washed once with cold (4 °C) PBS, and fixed with room temperature 4% PFA for 10 minutes. Neurons were incubated for 1 hour at room temperature with secondary antibodies to NMDAR subunit primary antibodies (Anti mouse IgG2a 594 or Anti rabbit 594), and then permeabilised with 0.1% Triton X-100 in PBS for 5 minutes. After blocking for 30 minutes with room temperature 3% BSA in PBS, neurons were incubated with primary antibodies to βIII-Tubulin or Tuj1 (Rabbit polyclonal or Mouse monoclonal, both 1:1000) and PSD95 (1:500) for 2 hours at room temperature. Finally, secondary antibodies to βIII-Tubulin and PSD95 (Anti rabbit 647 or Anti mouse 647 IgG2a and Anti mouse IgG1 488, respectively) were added for 1 hour at room temperature. After staining, the glass bottoms of the multiwell plates in which the neurons were grown were excised using a diamond tip pen and mounted on glass slides using Fluoroshield mounting medium (Sigma-Aldrich).

#### Structured Illumination Microscopy image acquisition and analysis

Neurons were imaged from three independent cultures per genotype. Imaging and analysis was carried out blind to genotype. 3D Structured Illumination Microscopy (SIM) image stacks were acquired using a Nikon N-SIM super resolution microscope equipped with an Apo TIRF 100x oil immersion objective (NA 1.49) and an Andor DU-897 EMCCD camera. The camera settings were as follows, and were kept consistent throughout the experiment. Readout speed: 1 MHz, bit depth: 16 bit, EM gain multiplier: 300, conversion gain: 1x, binning: 1×1. Laser power and/or exposure times were adjusted for each acquisition to produce the best signal to noise ratio, but keeping the grey levels in each image below 15,000. Each 3D-SIM stack consisted of 3 channels (green for PSD95, red for NMDAR subunits and far-red for βIII-Tubulin) and 17 slices, with the z-step size fixed at 120nm, for a total stack thickness of 2.04μm. To acquire each stack, the best focal plane was identified for each dendrite segment to be imaged, and 8 additional slices were acquired both above and below this central focal plane. Dendrites with clearly distinguishable morphological features (main shaft, primary branches and secondary branches), and no spatial overlap with other dendrites from neighbouring neurons, were selected for image acquisition. For each individual neuron analysed, a single dendrite segment measuring between 32 and 46μm in length and located immediately after the first dendrite branching point was imaged. The image reconstruction parameters (Illumination Modulation Contrast (IMC), High Resolution Noise Suppression (HRNS) and Out of Focus Blur Suppression (OFBS) were empirically selected for each channel to generate the best quality images, with minimal artefacts and the best Fourier transforms. The following reconstruction parameters were applied, and kept consistent throughout the experiment. Green channel (PSD95): IMC 0.3, HRNS 0.1, OFBS 0.05; Red channel (NMDAR subunit): IMC 0.5, HRNS 0.1, OFBS 0.05; Far-red channel (βIII-Tubulin): IMC 1, HRNS 5, OFBS 0.05. Lateral image resolution was determined by measuring the full-width half maximum (FWHM) of the smallest punctate structures detectable in a single plane of the reconstructed image, and was equal to 120 nm in the green channel (PSD95) and 180 nm in the red channel (NMDAR subunits).

Images were first analysed automatically using the Surfaces function of Imaris 8.1 and 8.4 (Bitplane). 3D segmentation parameters were first tested extensively for each channel on a sample of images acquired over different imaging sessions, and visually validated by superimposing the resulting 3D reconstruction with the raw images to confirm that all structures visible in the latter were faithfully represented in the former. The optimised parameters were then applied to the analysis of the entire dataset. The measured parameters included the volume of the imaged dendrite segment (estimated from the 3D rendering of βIII-Tubulin staining), as well as the number and volume of each individual punctum, and total volume of surface GluN1, GluN2A, GluN2B and of PSD95 proteins, based on the 3D reconstruction of their respective staining. Imaris 8.1 was also used to visualise and measure 3D co-localisation between each NMDAR subunit and PSD95 puncta. For intensity-based 3D co-localisation coefficients (Pearson’s and Mander’s), the above mentioned segmentation parameters were applied to each image to select the signal corresponding to either one of the NMDAR subunits and PSD95 puncta, and the intensity of the excluded voxels was considered below threshold and therefore set to zero.

#### Statistical analysis

GraphPad Prism software was used for statistical analyses. Pairwise comparisons were assessed using t-tests. Mutant neuron trafficking data were analysed using two-tailed t-tests, paired for each timepoint, to examine whether fluorescence intensities differed overall between genotypes within each bin. DISC1 overexpression trafficking data were analysed using the Friedman repeated measures test with post-hoc Dunn’s testing. Otherwise, multiple comparisons were carried out using one-way ANOVA with post-hoc pairwise Bonferroni tests or, if data were not normally distributed, by Kruskal-Wallis with post-hoc pairwise Dunn’s tests, as stated for each figure. Interaction between genotype and the number of PSD95 nanodomains per cluster was tested using two-way ANOVA with post-hoc Bonferroni testing. The hypergeometric probability test was used to examine gene enrichment in the RNASeq dataset. Statistical outliers, defined as being more than three standard deviations from the mean, were removed in one round from the GluN1-Dendra2 trafficking data and the surface expression data. All values are presented as mean ± SEM.

**Supplementary Figure 1.**
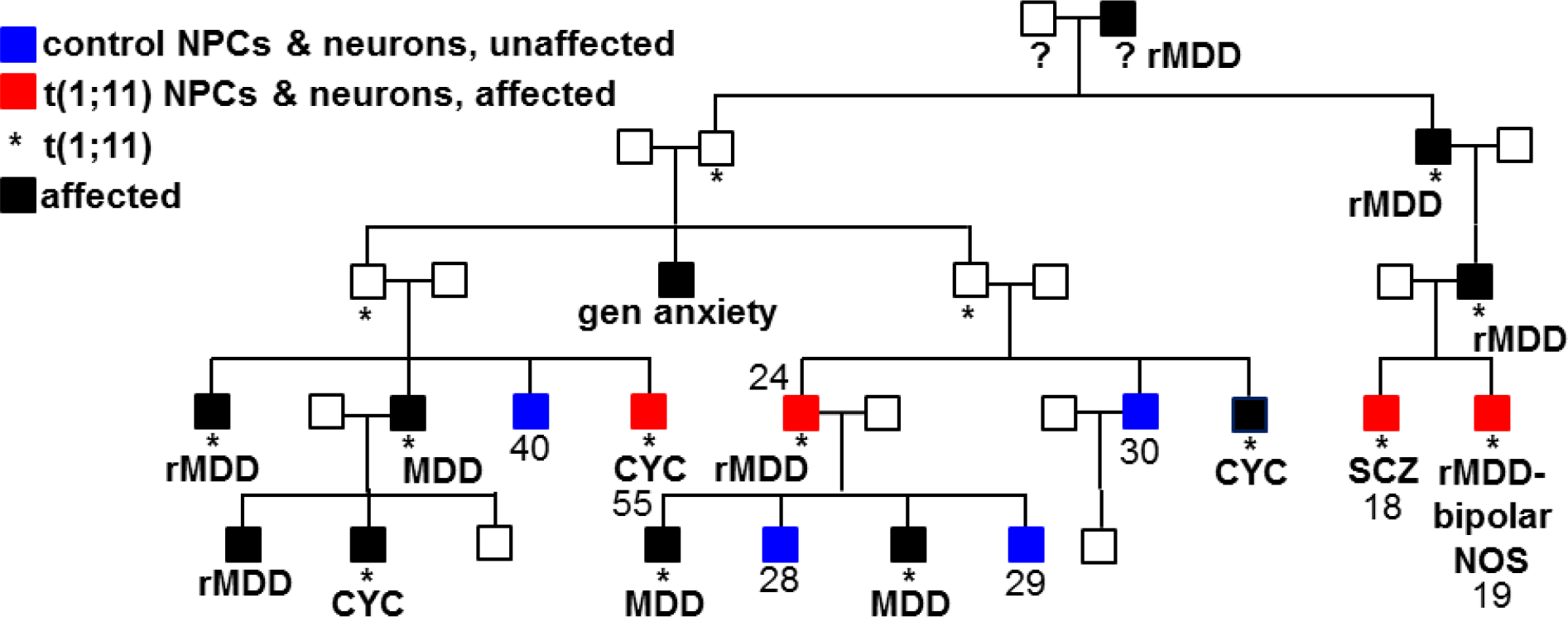
Translocation family pedigree indicating members from whom neural precursors and neurons were studied. rMDD, recurrent major depressive disorder; MDD, major depressive disorder; SCZ, schizophrenia; bipolar-NOS, bipolar disorder not otherwise specified; CYC, cyclothymia; ?, translocation carrier status unknown

**Supplementary Figure 2.**
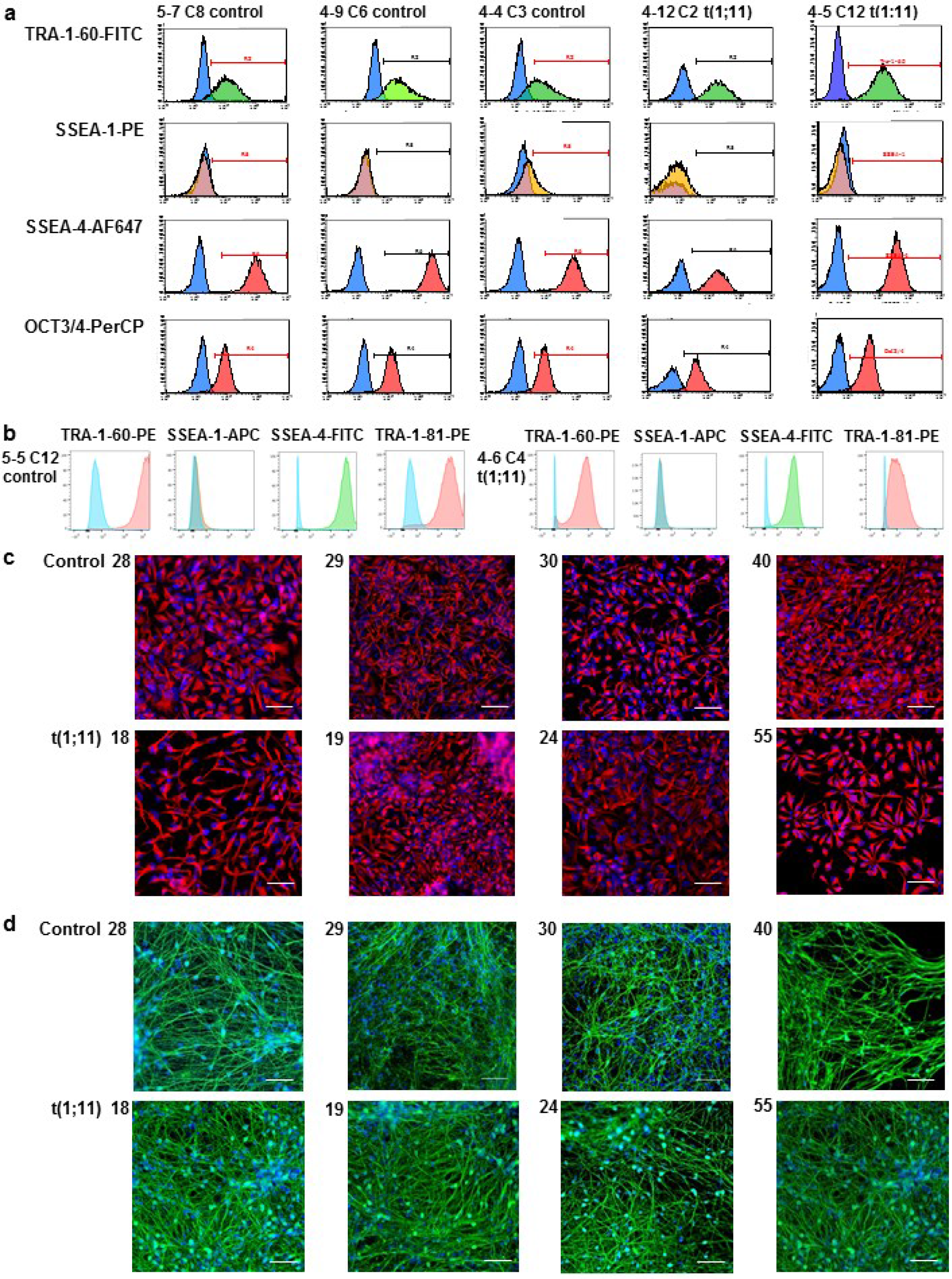
IPSC and neural cell controls. (**a**)(**b**) IPSC lines were subjected to flow cytometry using fluorescently labelled antibodies specific for cell surface markers. Flow cytometry detected the pluripotent stem cell markers TRA-1-60, SSEA-4 and OCT3/4, orTRA-1-60, SSEA-4 and TRA-1-81, respectively, but not the differentiation marker SSEA-1. y-axes, cell counts; x-axes, fluorescence; blue peaks, immunoglobulin isotype controls. (**c**) Representative examples of NPC lines stained for the neural stem/progenitor cell marker NES (red), with nuclei visualised using DAPI (blue). (**d**) Representative examples of neurons generated by differentiating NPC lines for five weeks, stained for the neuronal marker βIII-tubulin (green), with nuclei visualised using DAPI (blue). Scale bars, 5μm

**Supplementary Figure 3.**
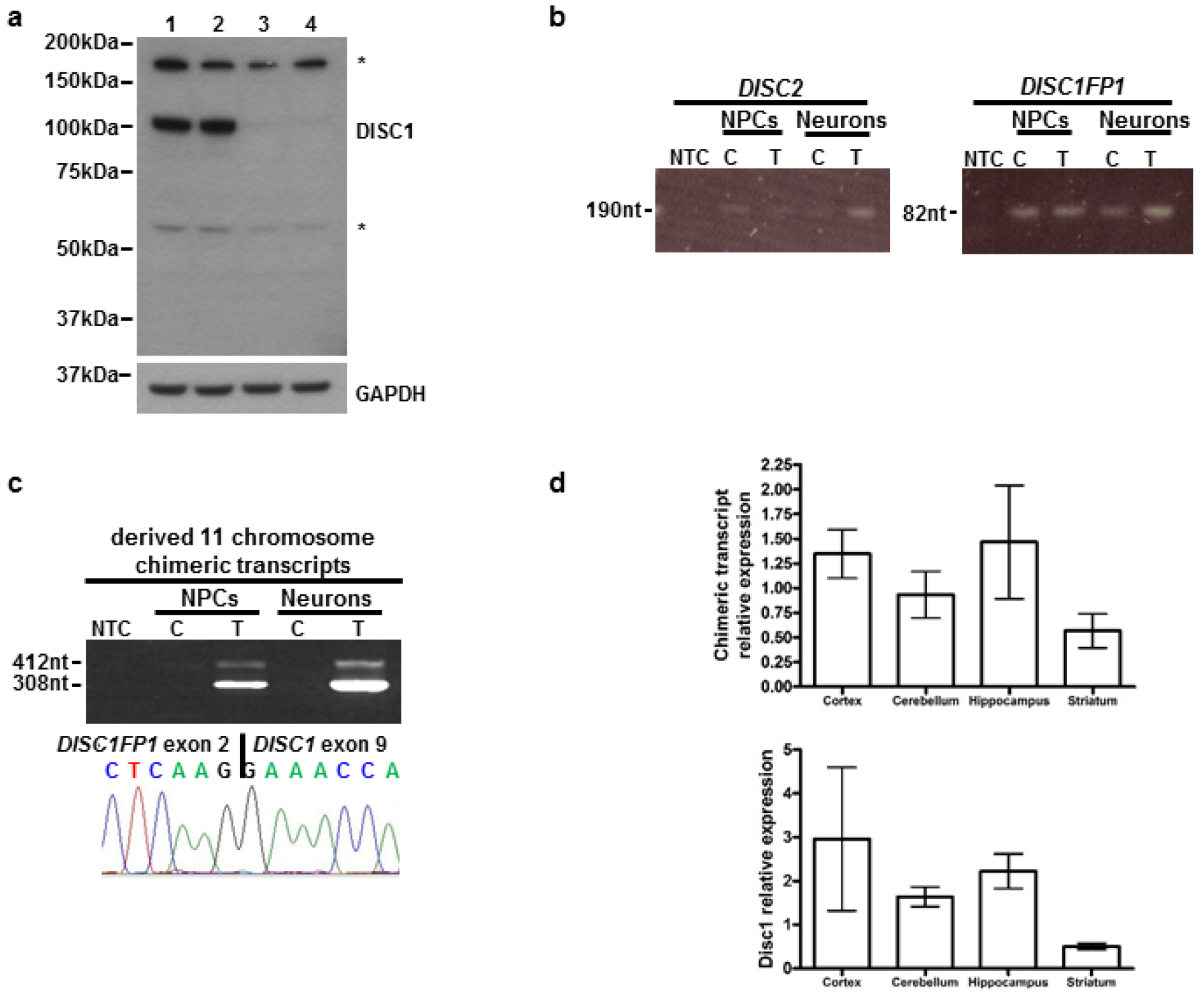
An antibody to detect human DISC1, and additional data from human NPCs & neurons, and from the *Der1* mouse. (**a**) Protein lysates from HEK293 cells transfected with previously tested human DISC1-specific siRNA duplexes ^9^, were immunoblotted and probed to demonstrate the specificity of a new antibody for full-length DISC1, or with GAPDH antibody. Only one species, at approximately 100kDa, is reduced by siRNA treatment. 1, mock transfection (no siRNA); 2, control siRNA; 3, DISC1-siRNA #2; 4, DISC1-siRNA #5; * non-specific bands. (**b**) Detection of *DISC2* and *DISC1FP1* expression by RT-PCR in neural precursors (NPCs) and neurons. Amplification specificity was confirmed by PCR product sequencing (data not shown). Expression of both genes was too low to be accurately quantified in these cell types. (**c**) Detection of chimeric transcripts from the derived 11 chromosome by RT-PCR followed by sequencing of the major product. Note that there are two products due to alternative splicing ^6^. (**d**) Quantification of chimeric (upper) or wild-type *Disc1* (lower) transcript expression in *Disc1*^*Der1/Der1*^ or *Disc1*^*wt/wt*^ adult mouse brain regions, respectively. N=4 per brain region per genotype; NTC, non-template control; C, control with normal karyotype; T, translocation carrier; error bars represent SEM

**Supplementary Figure 4.**
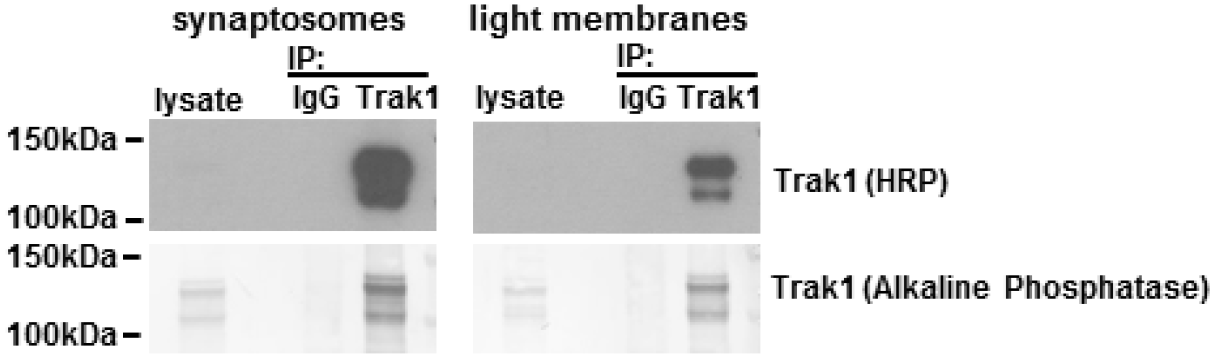
Trak1/GluN2B co-immunoprecipitation. Trak1 was immunoprecipitated from adult mouse brain synaptosome and light membrane fractions as shown in **Figure 3k**. Trak1 was detected in the lysates using secondary antibody conjugated to alkaline phosphatase.

**Supplementary Figure 5.**
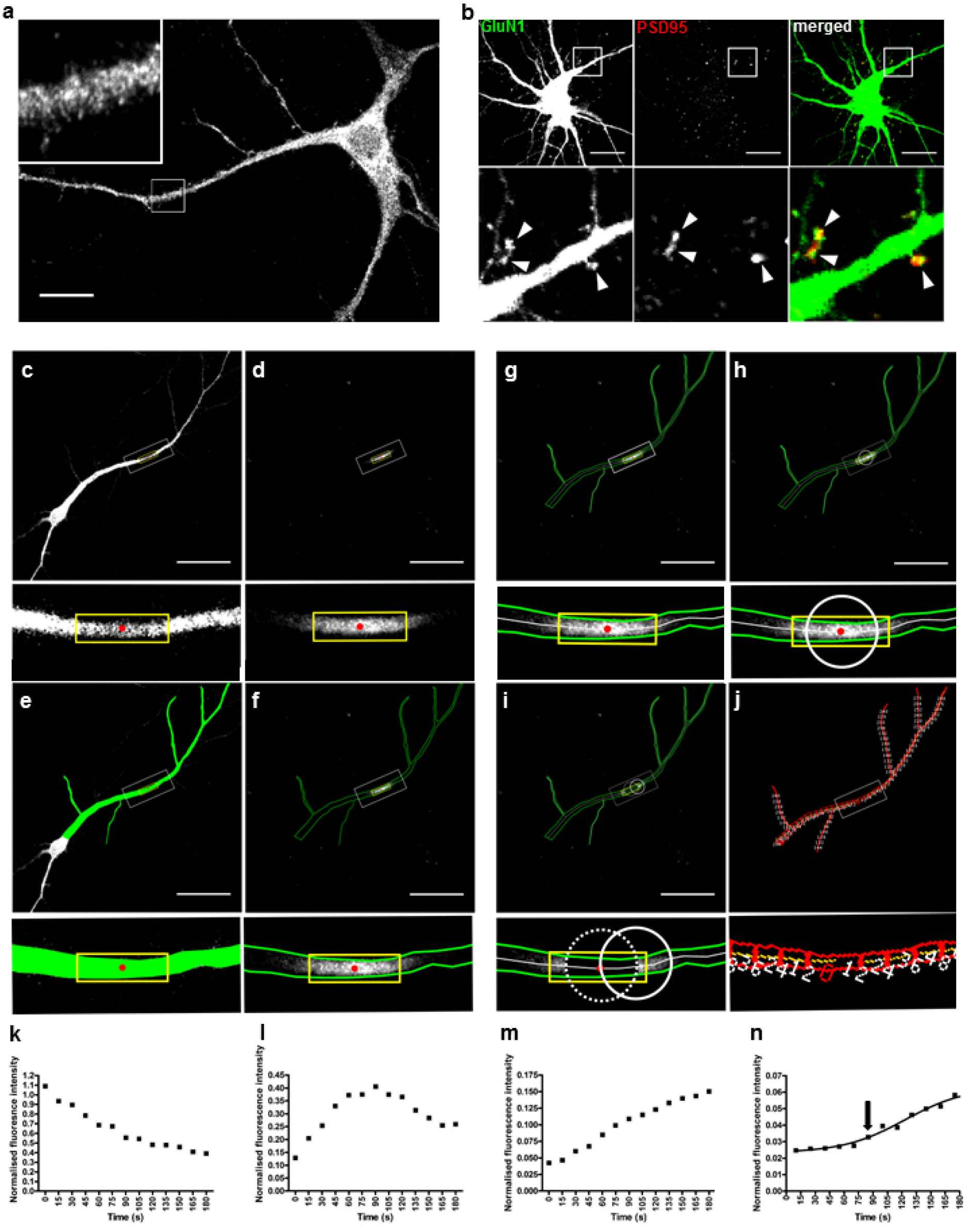
NMDAR trafficking assay controls and analysis of photoconverted GluN1-Dendra2 movement along a dendrite. (**a**) Endogenous GluN1 detected in wild-type DIV8 hippocampal neurons by immunofluorescence exhibits a fine granular appearance. scale bar, 15μm (**b**) GluN1-Dendra2 (native green Dendra2 fluorescence) co-localises with endogenous PSD95 in DIV14 hippocampal neurons within dendritic spines. scale bar, 20μm, arrowheads point to synapses (**c**)(**d**) Green fluorescent (non photoconverted) and red fluorescent (photoconverted) GluN1-Dendra2 in the first image frame captured after photoconversion. scale bars in c-j, 50μm (**e**) The dendrite area to be analysed is defined by a segmentation mask (green) applied on the green channel. (**f**) The area delimited by the segmentation mask is shown as a green line in the corresponding red channel image. Only the intensity of the red pixels located within the segmentation mask is measured. (**g**) A ‘skeleton’ (white line) corresponding to the longitudinal axis of the primary dendrite and all its branches is superimposed on the red image, providing a guide along which the algorithm will quantify the intensity of the red pixels. (**h**) At the start of the analysis, the analysis circle (white circle, 10μm in diameter) is placed on the geometric centre of the photoconversion ROI. Red pixels delimited by the intersection of the measuring circle with the dendrite outline (green) are measured through the time series. (**i**) The analysis circle moves one step along the dendrite in the distal direction. The new position of the centre of the analysis circle is defined by the intersection point between its previous position (dotted circle) and the dendrite skeleton (white line). The previous position of the circle (dotted circle) has been measured, and the corresponding data have been deleted. (**j**) Once the measuring circle has covered the entire dendritic area included in the image, an image showing all generated segments and their distance (in pixels) from the geometric centre of the photoconversion ROI is displayed. Mean intensity measured in all subsequent segments is normalised to the mean intensity measured in this first segment. (**k**) Example fluorescence intensity plot from a 5μm bin where fluorescence peaked within the 10 seconds prior to imaging onset. (**l**) 15μm bin where fluorescence peaked within the imaging period. (**m**) 25μm bin where fluorescence increased steadily because the peak was not reached during the imaging period. (**n**) 25μm bin where fluorescence appearance above background was delayed for 90s. This plot illustrates the use of sigmoidal curves to identify when fluorescence appears, as indicated by the arrow. Yellow rectangles represent the photoconversion ROI (14μm wide on average); red dots indicate the geometric centre of the photoconversion ROI; white rectangles indicate enlarged areas

**Figure 6.**
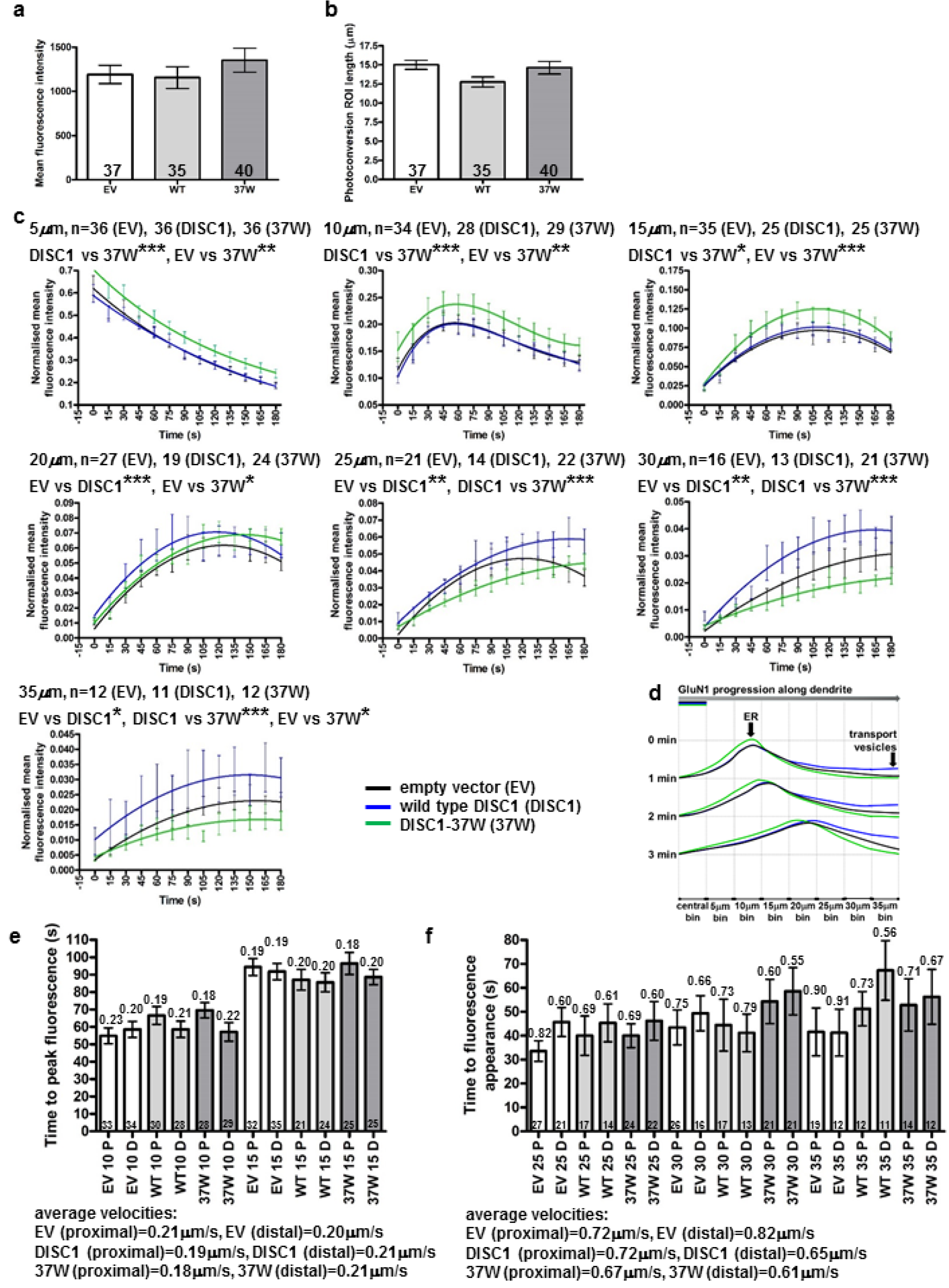
Altered distal dendritic NMDAR trafficking in mouse hippocampal neurons overexpressing DISC1 or DISC1-37W. (**a**)(**b**) Mean red fluorescence intensity, b, or ROI dendritic length, c, in the central bin in *Disc1*^*wt/wt*^ and *Disc1*^*Der1/Der1*^ DIV8 neurons was equal at time zero following photoconversion. (**c**) Quantification of fluorescence intensity over time in successive 5μm dendritic bins distal to the centre of the photoconversion ROI. Data analysed by timepoint-paired two tailed t-test. (**d**) Model of dendritic GluN1-Dendra2 motility. Photoconverted GluN1-Dendra2 progresses in a wave-like fashion, with the fastest and slowest moving GluN1-Dendra2 at the leading and trailing edges, respectively, and the bulk travelling as the ‘crest’. (**e**) Fluorescence peak velocity estimates for the 10μm and 15μm bins. Average time to peak fluorescence was converted to velocity, indicated above each bar. Average velocities were determined from the two bins. (**f**) Fast-moving GluN1-Dendra2 maximum velocity estimates for the 25μm-40μm bins. Average time to fluorescence appearance was converted to velocity, indicated above each bar. Average velocities were determined from the four bins. error bars represent SEM; **** p<0.0001; *** p<0.001; ** p<0.01; *p<0.05; n indicated on graphs

**Supplementary Figure 7.**
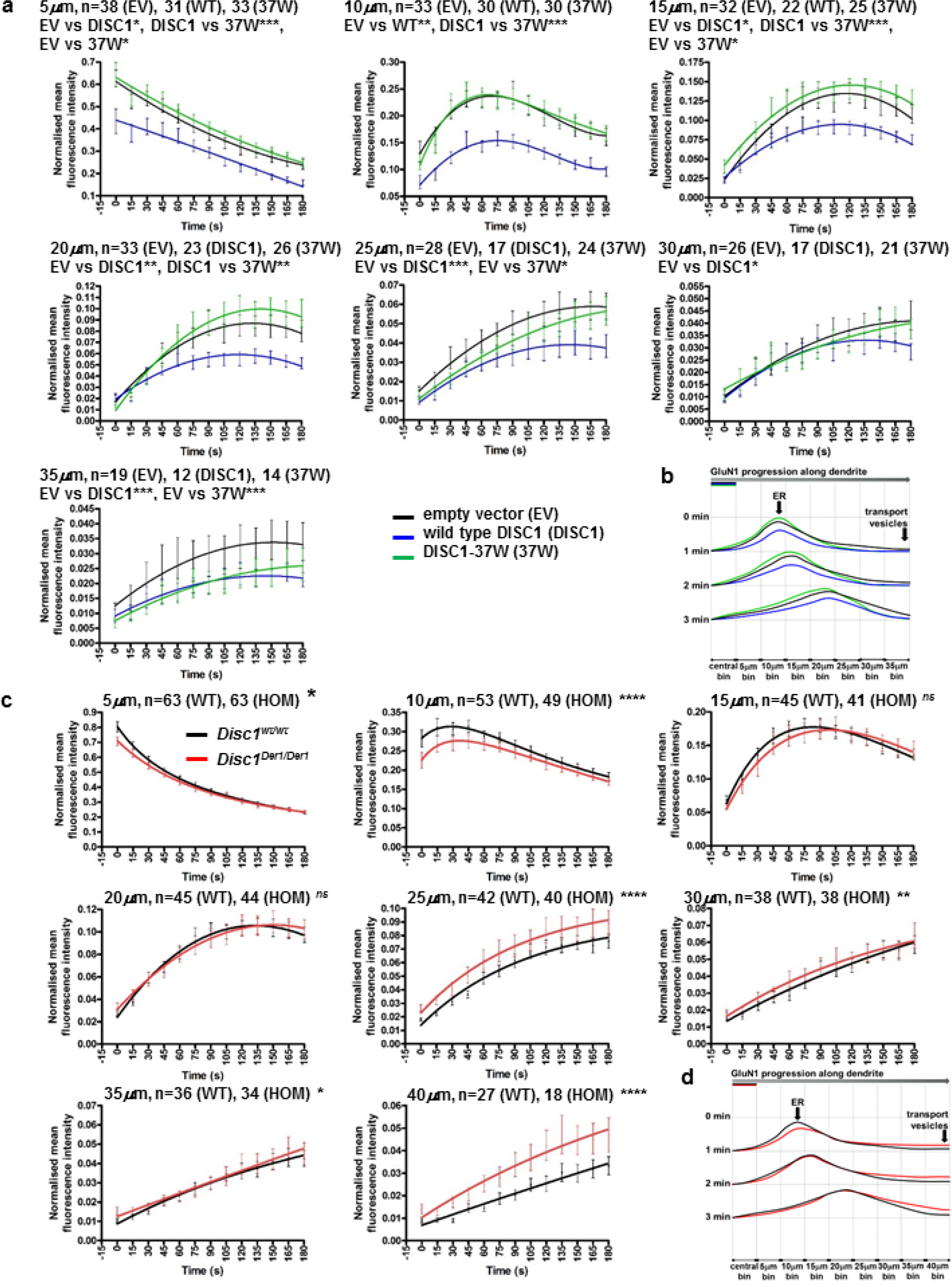
Altered proximal dendritic GluN1-Dendra2 motility due to DISC1 overexpression or the *Der1* mutation. (**a**) Effect of DISC1 overexpression. Quantification of fluorescence intensity over time in successive 5μm dendritic bins proximal to the centre of the photoconversion ROI. Mean fluorescence intensity at each time point within each bin is normalised to mean fluorescence intensity in the central bin at time zero per neuron. Total neuron numbers from three independent cultures are indicated. To determine whether fluorescence intensity differed between the expression constructs within each bin, data were analysed by Friedman repeated measures test (p<0.0001, p<0.0001, p<0.0001, p=0.0008, p<0.0001, p=0.02, p<0.0001, for the 5μm to 35μm bins, respectively) with Dunn’s post-hoc testing. (**b**) Model of dendritic GluN1-Dendra2 motility as described in **Figure 4e**. Total neuron numbers from three independent cultures are indicated. EV, empty vector; DISC1, wild-type DISC1; 37W, DISC1-37W; error bars represent SEM; *** p<0.001; ** p<0.01; *p<0.05 (**c**) Effect of the *Der1* mutation. Quantification of fluorescence intensity over time in successive 5μm bins proximal to the centre of the photoconversion ROI. Mean fluorescence intensity at each time point within each bin is normalised to mean fluorescence intensity in the central bin at time zero per neuron. Total neuron numbers from three independent cultures are indicated. To determine whether fluorescence intensity differed between the genotypes within each bin, data were analysed by paired two tailed t-test to make comparisons at each timepoint. (**d**) Model of dendritic GluN1-Dendra2 motility as described in **Figure 4e**. WT, *Disc1*^*wt/wt*^; HOM, *Disc1*^*Der1/Der1*^; error bars represent SEM, **** p<0.0001, ** p<0.01, * p<0.05, *ns* not significant

**Supplementary Figure 8.**
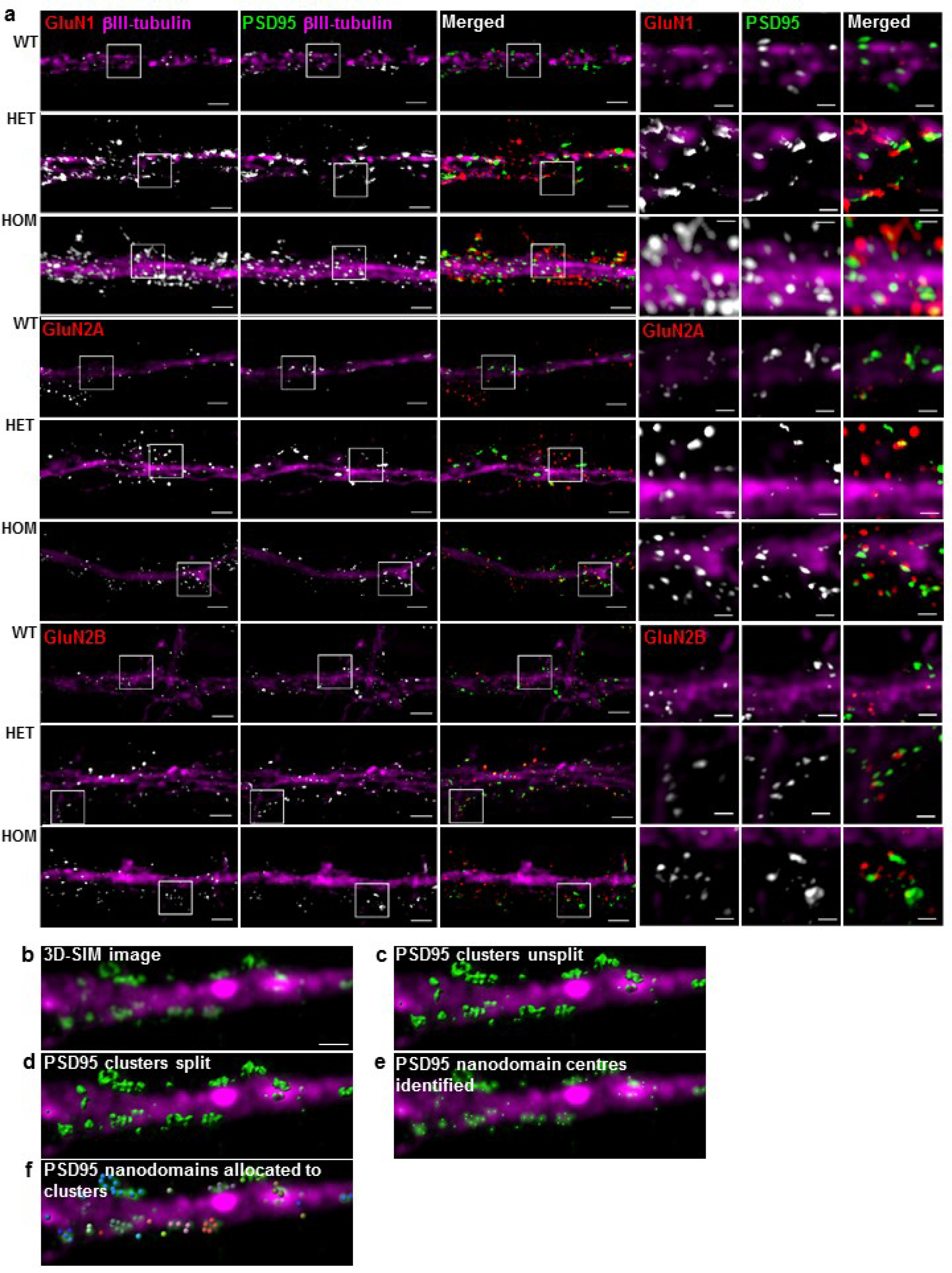
Dendritic NMDAR and PSD95 expression in hippocampal neurons. (**a**) 3D-SIM images of surface GluN1, GluN2A or Glun2B, and total PSD95 and βIII-tubulin (Tuj1). WT, *Disc1*^*wt/wt*^; HET, *Disc1*^*(wt)/Der1*^; HOM, *Disc1*^*Der1/Der1*^; scale bars, 2μm in main images, 0.6μm in enlarged insets indicated by white boxes (**b**) 3D-SIM image of a dendrite segment showing PSD95 (green) and βIII-tubulin (Tuj1, magenta). Scale bar B-F, 1μm (**c**) Identification of Imaris surfaces for PSD95. These three-dimensional surfaces are counted and their volume is quantified by the software. (**d**) PSD95 surfaces split into individual nanodomains. (**e**) Identification of the centre of each individual nanodomain, and conversion to Imaris spots using a bespoke MATLAB (MathWorks) XTension script. (**f**) Individual nanodomains are assigned to clusters, and the number per cluster is counted using the Imaris spots MATLAB XTension ‘Split into Surface Objects’.

**Supplementary Figure 9.**
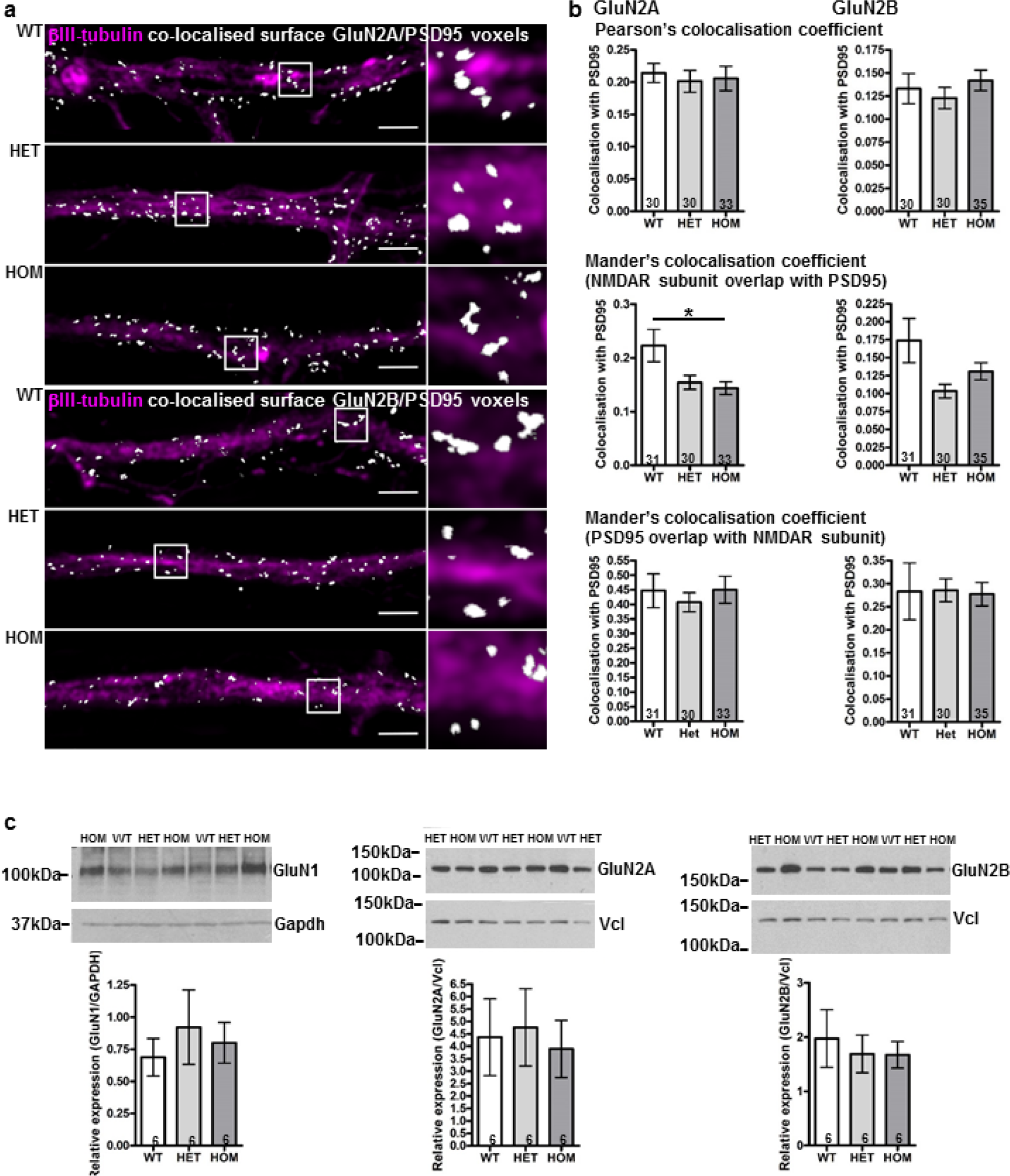
NMDAR subunit GluN2A and GluN2B co-localisation with the post-synaptic density marker PSD95. (**a**) Reconstructed 3D-SIM images of dendrites. Colocalised voxels, which contain signal from both PSD95 and surface-expressed GluN1 are shown in white. White boxes indicate enlarged regions. Scale bars, 3μm (**b**) Pearson’s coefficient indicates overall correlation of each signal. Mander’s coefficients measure the amount of subunit fluorescent signal co-localised with total PSD95 signal, and vice versa. Data were analysed by Kruskal-Wallis (p=0.02 for GluN2A Mander’s M1, p=0.06 for GluN2B Mander’s M1) followed by Dunn’s multiple comparison test. WT, *Disc1*^*wt/wt*^; HET, *Disc1*^*(wt)/Der1*^; HOM, *Disc1*^*Der1/Der1*^; error bars represent SEM; * p<0.05 (**c**) Immunoblots of hippocampus lysates were prepared from nine week mice, and probed with antibodies specific for GluN1. GluN2A and GluN2B NMDAR subunits, followed by loading controls Gapdh and Vcl. Subunit expression relative to the loading controls was quantified using densitometry. Data were analysed by Kruskal-Wallis test and no significant differences were found WT, *Disc1*^*wt/wt*^; HET, *Disc1*^*(wt)/Der(1)*^; HOM, *Disc1*^*Der(1)/Der(1)*^; error bars represent SEM; n indicated on graphs

**Supplementary Figure 10.**
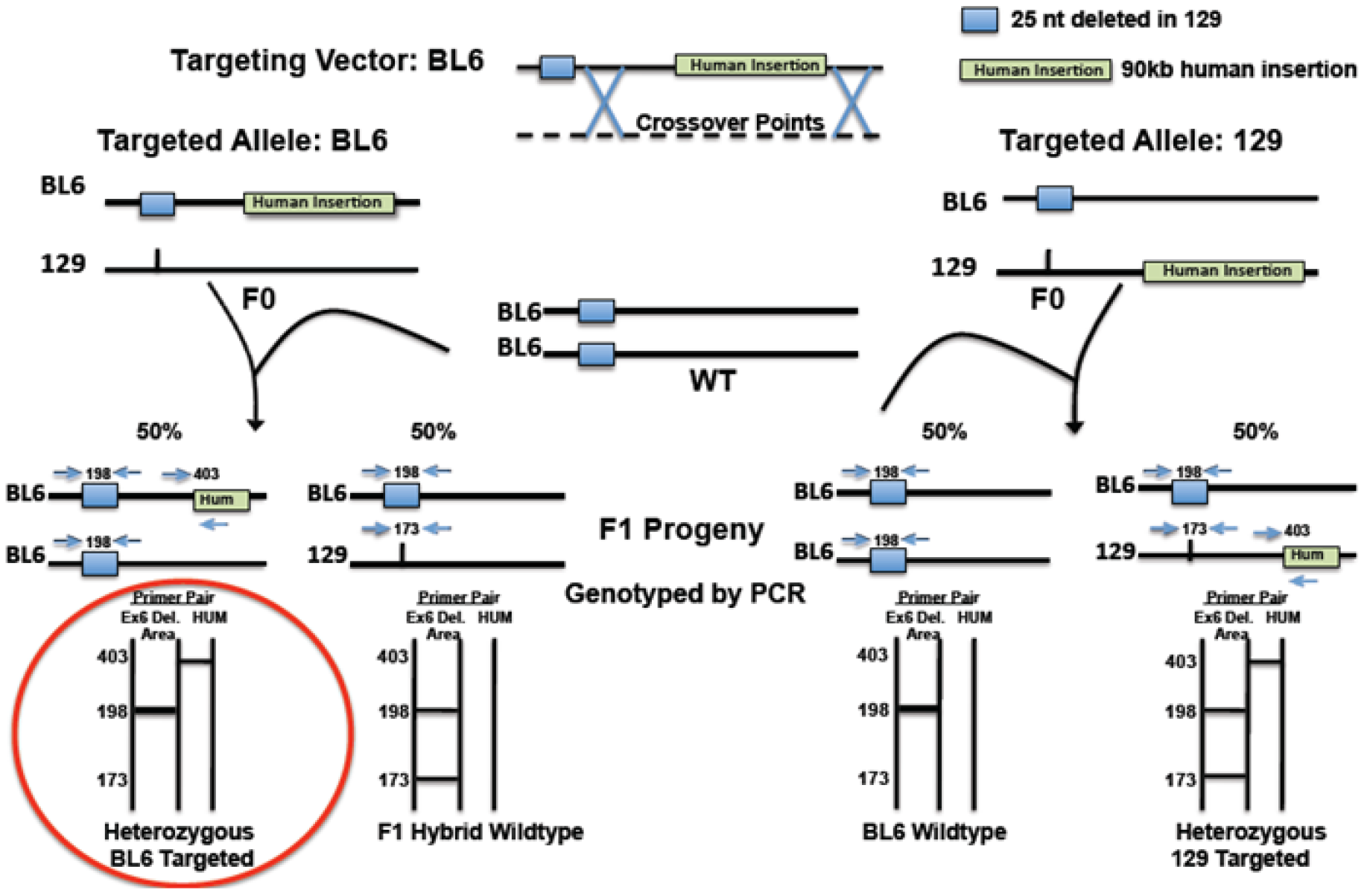
Screening for C57BL/6 targeted clones. PCR flanking the area of the naturally occurring *Disc1* deletion (25 base pairs)^5^ in strain 129S6SvEv distinguishes allele targeting. F1 progenies derived from C57BL/6 targeted clones produce a PCR product of 198nt from exon 6 and a 403 nt product from the human insertion.

**Supplementary Table 2.** Imaris image analysis data. Numbers of NMDAR subunit and PSD95 puncta counted for each genotype plus corresponding dendritic volumes. WT, *Disc1*^*wt/wt*^; HET, *Disc1*^*wt/Der1*^; HOM, *Disc1*^*Der1/Der1*^

|  | WT | HET | HOM |
| --- | --- | --- | --- |
| **GluN1** | 5,032 | 7,579 | 12,477 |
| **dendritic volume (μm<sup>3</sup>)** | 3,773 | 2,740 | 4,259 |
| **GluN2A** | 6,428 | 6,158 | 7,618 |
| **dendritic volume (μm<sup>3</sup>)** | 4,584 | 3,687 | 3,628 |
| **GluN2B** | 6,522 | 6,769 | 10,959 |
| **dendritic volume (μm<sup>3</sup>)** | 4,523 | 2,805 | 5,336 |
| **PSD95 unsplit** | 14,669 | 14,025 | 20,953 |
| **PSD95 split** | 23,306 | 22,775 | 33,170 |
| **total dendritic volume (μm<sup>3</sup>)** | 12,881 | 9,232 | 13,223 |

## REFERENCES

1. Howes O, McCutcheon R, Stone J. Glutamate and dopamine in schizophrenia: an update for the 21st century. J Psychopharmacol. 2015;29:97–115.

2. Schizophrenia Working Group of the Psychiatric Genomics Consortium. Biological insights from 108 schizophrenia-associated genetic loci. Nature 2014;511:421–427.

3. CNV, Schizophrenia Working Groups of the Psychiatric Genomics Consortium, Psychosis Endophenotypes International Consortium. Contribution of copy number variants to schizophrenia from a genome-wide study of 41,321 subjects. Nature Genet. 2017;49:27–35.

4. Hall J, Trent S, Thomas KL, O’Donovan MC, Owen MJ. Genetic risk for schizophrenia: convergence on synaptic pathways involved in plasticity. Biol Psychiatry. 2015;77:52–58.

5. Horak M, Petralia RS, Kaniakova M, Sans N. ER to synapse trafficking of NMDA receptors. Front Cell Neurosci. 2014;8:394.

6. Blackwood DH, Fordyce A, Walker MT, St Clair DM, Porteous DJ, Muir WJ. Schizophrenia and affective disorders--cosegregation with a translocation at chromosome 1q42 that directly disrupts brain-expressed genes: clinical and P300 findings in a family. Am J Hum Genet. 2001;69:428–433.

7. Millar JK, Wilson-Annan JC, Anderson S, Christie S, Taylor MS, Semple CA et al. Disruption of two novel genes by a translocation co-segregating with schizophrenia. Hum Mol Genet. 2000;9:1415–1423.

8. Thomson PA, Duff B, Blackwood DH, Romaniuk L, Watson A, Whalley HC et al. Balanced translocation linked to psychiatric disorder, glutamate, and cortical structure/function. NPJ Schizophr. 2016;2:16024.

9. Brandon NJ, Sawa A. Linking neurodevelopmental and synaptic theories of mental illness through DISC1. Nature Rev 2011;12:707–722.

10. Thomson PA, Malavasi EL, Grunewald E, Soares DC, Borkowska M, Millar JK. DISC1 genetics, biology and psychiatric illness. Front Biol. 2013;8:1–31.

11. Hayashi-Takagi A, Takaki M, Graziane N, Seshadri S, Murdoch H, Dunlop AJ et al. Disrupted-in-Schizophrenia 1 (DISC1) regulates spines of the glutamate synapse via Rac1. Nature Neurosci. 2010;13:327–332.

12. Wei J, Graziane NM, Wang H, Zhong P, Wang Q, Liu W et al. Regulation of N-Methyl-D-Aspartate Receptors by Disrupted-in-Schizophrenia-1. Biol Psychiatry. 2013;75:414–424

13. Flores R, 3rd, Hirota Y, Armstrong B, Sawa A, Tomoda T. DISC1 regulates synaptic vesicle transport via a lithium-sensitive pathway. Neurosci Res. 2011;71:71–77.

14. Murphy LC, Millar JK. Regulation of mitochondrial dynamics by DISC1, a putative risk factor for major mental illness. Schizophr Res. 2017;187:55–61

15. Tsuboi D, Kuroda K, Tanaka M, Namba T, Iizuka Y, Taya S et al. Disrupted-in-schizophrenia 1 regulates transport of ITPR1 mRNA for synaptic plasticity. Nature Neurosci. 2015;18:698–707.

16. Norkett R, Modi S, Birsa N, Atkin TA, Ivankovic D, Pathania M et al. DISC1-dependent Regulation of Mitochondrial Dynamics Controls the Morphogenesis of Complex Neuronal Dendrites. J Biol Chem. 2016;291:613–629.

17. Ogawa F, Malavasi EL, Crummie DK, Eykelenboom JE, Soares DC, Mackie S et al. DISC1 complexes with TRAK1 and Miro1 to modulate anterograde axonal mitochondrial trafficking. Hum Mol Genet. 2014;23:906–919.

18. Okita K, Matsumura Y, Sato Y, Okada A, Morizane A, Okamoto S et al. A more efficient method to generate integration-free human iPS cells. Nat Methods. 2011;8:409–412.

19. Bilican B, Livesey MR, Haghi G, Qiu J, Burr K, Siller R et al. Physiological normoxia and absence of EGF is required for the long-term propagation of anterior neural precursors from human pluripotent cells. PLoS One. 2014;9:e85932

20. Darmanis S, Sloan SA, Zhang Y, Enge M, Caneda C, Shuer LM et al. A survey of human brain transcriptome diversity at the single cell level. Proc Natl Acad Sci USA. 2015;112:7285–7290.

21. Zhang Y, Chen K, Sloan SA, Bennett ML, Scholze AR, O’Keeffe S et al. An RNA-sequencing transcriptome and splicing database of glia, neurons, and vascular cells of the cerebral cortex. J Neurosci. 2014;34:11929–11947.

22. Srikanth P, Han K, Callahan DG, Makovkina E, Muratore CR, Lalli MA et al. Genomic DISC1 Disruption in hiPSCs Alters Wnt Signaling and Neural Cell Fate. Cell Rep. 2015;12:1414–1429.

23. Wen Z, Nguyen HN, Guo Z, Lalli MA, Wang X, Su Y et al. Synaptic dysregulation in a human iPS cell model of mental disorders. Nature. 2014;515:414–418.

24. Pardinas AF, Holmans P, Pocklington AJ, Escott-Price V, Ripke S, Carrera N et al. Common schizophrenia alleles are enriched in mutation-intolerant genes and in regions under strong background selection. Nature Genet. 2018;50:381–389.

25. Millar JK, Pickard BS, Mackie S, James R, Christie S, Buchanan SR et al. DISC1 and PDE4B are interacting genetic factors in schizophrenia that regulate cAMP signaling. Science. 2005;310:1187–1191.

26. Mostaid MS, Lloyd D, Liberg B, Sundram S, Pereira A, Pantelis C et al. Neuregulin-1 and schizophrenia in the genome-wide association study era. Neurosci Biobehav Rev. 2016;68:387–409.

27. Seshadri S, Faust T, Ishizuka K, Delevich K, Chung Y, Kim SH et al. Interneuronal DISC1 regulates NRG1-ErbB4 signalling and excitatory-inhibitory synapse formation in the mature cortex. Nat Commun. 2015;6:10118.

28. Unda BK, Kwan V, Singh KK. Neuregulin-1 Regulates Cortical Inhibitory Neuron Dendrite and Synapse Growth through DISC1. Neural Plast. 2016;2016:7694385.

29. Wray NR, Ripke S, Mattheisen M, Trzaskowski M, Byrne EM, Abdellaoui A et al. Genome-wide association analyses identify 44 risk variants and refine the genetic architecture of major depression. Nature Genet. 2018;50:668–681.

30. Ryan NM, Lihm J, Kramer M, McCarthy S, Morris SW, Arnau-Soler A et al. DNA sequence-level analyses reveal potential phenotypic modifiers in a large family with psychiatric disorders. Mol Psychiatry 2018;epub

31. Coba MP, Komiyama NH, Nithianantharajah J, Kopanitsa MV, Indersmitten T, Skene NG et al. TNiK is required for postsynaptic and nuclear signaling pathways and cognitive function. J Neurosci. 2012;32:13987–13999.

32. Das SR, Magnusson KR. Changes in expression of splice cassettes of NMDA receptor GluN1 subunits within the frontal lobe and memory in mice during aging. Behav Brain Res 2011;222:122–133.

33. Ogawa F, Murphy LC, Malavasi EL, O’Sullivan ST, Torrance HS, Porteous DJ et al. NDE1 and GSK3beta Associate with TRAK1 and Regulate Axonal Mitochondrial Motility: Identification of Cyclic AMP as a Novel Modulator of Axonal Mitochondrial Trafficking. ACS Chemical Neurosci. 2016;7:553–564.

34. Tropea D, Hardingham N, Millar K, Fox K. Mechanisms underlying the role of DISC1 in synaptic plasticity. J Physiol. (in press)

35. Gurskaya NG, Verkhusha VV, Shcheglov AS, Staroverov DB, Chepurnykh TV, Fradkov AF et al. Engineering of a monomeric green-to-red photoactivatable fluorescent protein induced by blue light. Nat Biotechnol. 2006;24:461–465.

36. Song W, Li W, Feng J, Heston LL, Scaringe WA, Sommer SS. Identification of high risk DISC1 structural variants with a 2% attributable risk for schizophrenia. Biochem Biophys Res Comm. 2008;367:700–706.

37. Thomson PA, Parla JS, McRae AF, Kramer M, Ramakrishnan K, Yao J et al. 708 Common and 2010 rare DISC1 locus variants identified in 1542 subjects: analysis for association with psychiatric disorder and cognitive traits. Mol Psychiatry. 2014;19:668–675.

38. Mironov SL, Symonchuk N. ER vesicles and mitochondria move and communicate at synapses. J Cell Sci 2006;119:4926–4934.

39. Guillaud L, Setou M, Hirokawa N. KIF17 dynamics and regulation of NR2B trafficking in hippocampal neurons. J Neurosci. 2003;23:131–140.

40. Kamiya A, Kubo K, Tomoda T, Takaki M, Youn R, Ozeki Y et al. A schizophrenia-associated mutation of DISC1 perturbs cerebral cortex development. Nature Cell Biol. 2005;7:1167–1178.

41. Sheng M, Kim E. The postsynaptic organization of synapses. Cold Spring Harb Perspect Biol. 2011;3.

42. Nair D, Hosy E, Petersen JD, Constals A, Giannone G, Choquet D et al. Superresolution imaging reveals that AMPA receptors inside synapses are dynamically organized in nanodomains regulated by PSD95. J Neurosci. 2013;33:13204–13224.

43. Fukata Y, Dimitrov A, Boncompain G, Vielemeyer O, Perez F, Fukata M. Local palmitoylation cycles define activity-regulated postsynaptic subdomains. J Cell Biol 2013;202:145–161.

44. Broadhead MJ, Horrocks MH, Zhu F, Muresan L, Benavides-Piccione R, DeFelipe J et al. PSD95 nanoclusters are postsynaptic building blocks in hippocampus circuits. Sci Rep. 2016;6:24626.

45. Newpher TM, Ehlers MD. Glutamate receptor dynamics in dendritic microdomains. Neuron. 2008;58:472–497.

46. Triller A, Choquet D. Surface trafficking of receptors between synaptic and extrasynaptic membranes: and yet they do move! Trends Neurosci. 2005;28:133–139.

47. Liu KK, Hagan MF, Lisman JE. Gradation (approx. 10 size states) of synaptic strength by quantal addition of structural modules. Philos Trans R Soc Lond B Biol Sci. 2017;372.

48. Havekes R, Park AJ, Tolentino RE, Bruinenberg VM, Tudor JC, Lee Y et al. Compartmentalized PDE4A5 Signaling Impairs Hippocampal Synaptic Plasticity and Long-Term Memory. J Neurosci. 2016;36:8936–8946.

49. Song RS, Massenburg B, Wenderski W, Jayaraman V, Thompson L, Neves SR. ERK regulation of phosphodiesterase 4 enhances dopamine-stimulated AMPA receptor membrane insertion. Proc Natl Acad Sci USA. 2013;110:15437–15442.

50. Vogel EW, 3rd, Morales FN, Meaney DF, Bass CR, Morrison B, 3rd. Phosphodiesterase-4 inhibition restored hippocampal long term potentiation after primary blast. Exp Neurol. 2017;293:91–100.

51. Wang H, Peng RY. Basic roles of key molecules connected with NMDAR signaling pathway on regulating learning and memory and synaptic plasticity. Mil Med Res. 2016;3:26.

52. Wiescholleck V, Manahan-Vaughan D. PDE4 inhibition enhances hippocampal synaptic plasticity in vivo and rescues MK801-induced impairment of long-term potentiation and object recognition memory in an animal model of psychosis. Transl Psychiatry. 2012;2:e89.

53. Bosch M, Castro J, Saneyoshi T, Matsuno H, Sur M, Hayashi Y. Structural and molecular remodeling of dendritic spine substructures during long-term potentiation. Neuron. 2014;82:444–459.

54. Herring BE, Nicoll RA. Long-Term Potentiation: From CaMKII to AMPA Receptor Trafficking. Annu Rev Physiol. 2016;78:351–365.

55. Wang Q, Charych EI, Pulito VL, Lee JB, Graziane NM, Crozier RA et al. The psychiatric disease risk factors DISC1 and TNIK interact to regulate synapse composition and function. Mol Psychiatry. 2011;16:1006–1023.

56. Koychev I, William Deakin JF, El-Deredy W, Haenschel C. Effects of Acute Ketamine Infusion on Visual Working Memory: Event-Related Potentials. Biol Psychiatry Cogn Neurosci Neuroimaging. 2017;2:253–262.

## SUPPLEMENTARY REFERENCES

1. Okita K, Matsumura Y, Sato Y, Okada A, Morizane A, Okamoto S et al. A more efficient method to generate integration-free human iPS cells. Nat Methods. 2011;8:409–412.

2. Chambers SM, Fasano CA, Papapetrou EP, Tomishima M, Sadelain M, Studer L. Highly efficient neural conversion of human ES and iPS cells by dual inhibition of SMAD signaling. Nat Biotechnol. 2009;27:275–280.

3. Bilican B, Livesey MR, Haghi G, Qiu J, Burr K, Siller R et al. Physiological normoxia and absence of EGF is required for the long-term propagation of anterior neural precursors from human pluripotent cells. PLoS One. 2014;9:e85932.

4. Dechiara TM, Poueymirou WT, Auerbach W, Frendewey D, Yancopoulos GD, Valenzuela DM. VelociMouse: fully ES cell-derived F0-generation mice obtained from the injection of ES cells into eight-cell-stage embryos. Methods Mol Biol. 2009;530:311–324.

5. Koike H, Arguello PA, Kvajo M, Karayiorgou M, Gogos JA. Disc1 is mutated in the 129S6/SvEv strain and modulates working memory in mice. Proc Natl Acad Sci USA. 2006;103:3693–3697.

6. Eykelenboom JE, Briggs GJ, Bradshaw NJ, Soares DC, Ogawa F, Christie S et al. A t(1;11) translocation linked to schizophrenia and affective disorders gives rise to aberrant chimeric DISC1 transcripts that encode structurally altered, deleterious mitochondrial proteins. Hum Mol Genet. 2012;21:3374–3386.

7. Jeyifous O, Waites CL, Specht CG, Fujisawa S, Schubert M, Lin EI et al. SAP97 and CASK mediate sorting of NMDA receptors through a previously unknown secretory pathway. Nature Neurosci. 2009;12:1011–1019.

8. Malavasi EL, Ogawa F, Porteous DJ, Millar JK. DISC1 variants 37W and 607F disrupt its nuclear targeting and regulatory role in ATF4-mediated transcription. Hum Mol Genet. 2012;21:2779–2792.

9. Ogawa F, Malavasi EL, Crummie DK, Eykelenboom JE, Soares DC, Mackie S et al. DISC1 complexes with TRAK1 and Miro1 to modulate anterograde axonal mitochondrial trafficking. Hum Mol Genet. 2014;23:906–919.

10. Love MI, Huber W, Anders S. Moderated estimation of fold change and dispersion for RNA-seq data with DESeq2. Genome Biol. 2014;15:550.

11. Anders S, Reyes A, Huber W. Detecting differential usage of exons from RNA-seq data. Genome Res. 2012;22:2008–2017.

12. Baillie GS, Adams DR, Bhari N, Houslay TM, Vadrevu S, Meng D et al. Mapping binding sites for the PDE4D5 cAMP-specific phosphodiesterase to the N-and C-domains of beta-arrestin using spot-immobilized peptide arrays. Biochem J. 2007;404:71–80.

13. Bolger GB, Baillie GS, Li X, Lynch MJ, Herzyk P, Mohamed A et al. Scanning peptide array analyses identify overlapping binding sites for the signalling scaffold proteins, beta-arrestin and RACK1, in cAMP-specific phosphodiesterase PDE4D5. Biochem J. 2006;398:23–36.

14. Clapcote SJ, Lipina TV, Millar JK, Mackie S, Christie S, Ogawa F et al. Behavioral phenotypes of Disc1 missense mutations in mice. Neuron. 2007;54: 87–402.

15. Ogawa F, Murphy LC, Malavasi EL, O’Sullivan ST, Torrance HS, Porteous DJ et al. NDE1 and GSK3beta Associate with TRAK1 and Regulate Axonal Mitochondrial Motility: Identification of Cyclic AMP as a Novel Modulator of Axonal Mitochondrial Trafficking. ACS ChemNeurosci. 2016;7:553–564.

16. Chandran JS, Kazanis I, Clapcote SJ, Ogawa F, Millar JK, Porteous DJ et al. Disc1 variation leads to specific alterations in adult neurogenesis. PLoS One. 2014;9:e108088.

17. Kaech S, Banker G. Culturing hippocampal neurons. Nature Prot. 2006;1:2406–2415.

18. Horak M, Petralia RS, Kaniakova M, Sans N. ER to synapse trafficking of NMDA receptors. Front Cell Neurosci. 2014;8:394.

